# Small molecule sequestration of amyloid-β as a drug discovery strategy for Alzheimer’s disease

**DOI:** 10.1101/729392

**Authors:** Gabriella T. Heller, Francesco A. Aprile, Thomas C. T. Michaels, Ryan Limbocker, Michele Perni, Francesco Simone Ruggeri, Benedetta Mannini, Thomas Löhr, Massimiliano Bonomi, Carlo Camilloni, Alfonso De Simone, Isabella C. Felli, Roberta Pierattelli, Tuomas P. J. Knowles, Christopher M. Dobson, Michele Vendruscolo

**Affiliations:** Centre for Misfolding Diseases, Department of Chemistry, University of Cambridge, Cambridge CB2 1EW, UK; Paulson School of Engineering and Applied Sciences, Harvard University, Cambridge MA, USA; Structural Bioinformatics Unit, Department of Structural Biology and Chemistry; CNRS UMR 3528; C3BI, CNRS USR 3756; Institut Pasteur, Paris, France; Dipartimento di Bioscienze, Università degli Studi di Milano, via Celoria 26, 20133 Milano,Italy; Division of Molecular Biosciences, Imperial College London, London SW7 2AZ, UK; Magnetic Resonance Center (CERM), University of Florence, 50019 Sesto Fiorentino, Italy; Department of Chemistry “Ugo Schiff”, University of Florence, 50019 Sesto Fiorentino, Italy

## Abstract

Disordered proteins are challenging therapeutic targets, and no drug is currently in clinical use that has been shown to modify the properties of their monomeric states. Here, we identify a small molecule capable of binding and sequestering the amyloid-β peptide (Aβ) in its monomeric, soluble state. Our analysis reveals that this compound interacts with Aβ and inhibits both the primary and secondary nucleation pathways in its aggregation process. We characterise this interaction using biophysical experiments and integrative structural ensemble determination methods. We thus observe that this small molecule has the remarkable effect of increasing the conformational entropy of monomeric Aβ while decreasing its hydrophobic surface area. We then show that this small molecule rescues a *Caenorhabditis elegans* model of Aβ-associated toxicity in a manner consistent with the mechanism of action identified from the *in silico* and *in vitro* studies. These results provide an illustration of the strategy of targeting the monomeric states of disordered proteins with small molecules to alter their behaviour for therapeutic purposes.

## Introduction

Alzheimer’s disease is a fatal neurodegenerative condition that affects over 50 million people worldwide, a number that is predicted to rise to 150 million by 2050 unless methods of prevention or treatment are found, with a cost to the global economy that exceeds one trillion dollars per year^1,2^. Despite over 25 years of intensive research and hundreds of clinical trials, there is still no drug capable of modifying the course of this disease^1,2^.

The aggregation of the amyloid-β peptide (Aβ) in brain tissue is one of the hallmarks of Alzheimer’s disease^3-7^. This process involves at least three forms of Aβ: (i) a monomeric state, which is highly disordered, (ii) oligomeric aggregates, which are heterogeneous, transient and cytotoxic, and (iii) fibrillar structures, which are ordered and relatively inert, although they are capable of catalysing the formation of Aβ oligomers^8,9^. More generally, the aggregation of Aβ involves a complex non-linear network of inter-dependent microscopic processes, including: (1) primary nucleation, in which oligomers form from monomeric species, (2) elongation, in which oligomers and fibrils increase in size by monomer addition, (3) secondary nucleation, whereby the surfaces of fibrillar aggregates catalyse the formation of new oligomeric species, and (4) fragmentation, in which fibrils break into smaller pieces, increasing the total number of oligomers and fibrils capable of elongation^10,11^.

Aβ is produced by proteolysis from the transmembrane amyloid precursor protein, and its 42-residue form (Aβ42) is the predominant species in deposits characteristically observed in the brains of patients with Alzheimer’s disease^6,7,12^. Kinetic analysis shows that, once a critical concentration of Aβ42 fibrils has been formed, secondary nucleation overtakes primary nucleation in becoming the major source of Aβ42 oligomers, as fibril surfaces act as catalytic sites for their formation^8^. The fact that the oligomers appear to be the most toxic species formed during the aggregation process^13-15^, however, suggests that therapeutic strategies targeting Aβ aggregation should not primarily aim at inhibiting fibril formation per se, but rather doing so in a manner that specifically reduces the generation of oligomeric species^16^. Complex feedback mechanisms between the different microscopic steps in the aggregation reaction can lead to an increase in the concentration of oligomers even when the formation of fibrils is inhibited, and hence result in an increase in pathogenicity^16^.

Previous studies have suggested that effective strategies for inhibiting Aβ aggregation could be based on targeting fibril surfaces to supress the generation of oligomers, or on the reduction of the toxicity of the oligomers^17-21^. It is unclear, however, whether sequestering Aβ in its soluble state could be an effective drug discovery strategy against Alzheimer’s disease. Stabilisation of monomeric Aβ into a β-hairpin conformation with large biomolecules has been previously demonstrated to inhibit aggregation, for example using an affibody protein^22^. However, whether such stabilisation of Aβ in its monomeric form can be achieved via small molecule binding in a drug-like manner is still under debate. While there is research indicating a stabilising effect of small molecules on the soluble state of Aβ, there are contradictory reports of their effects on its aggregation^23-25^. It should also be considered that such molecules may not be specific, as for example some appear to bind monomeric Aβ in a manner similar to low concentrations of sodium dodecyl sulphate (SDS)^23-25^. Furthermore, it has been proposed that the binding of these small molecules to monomeric Aβ may be mediated by colloidal particles formed by the small molecules^26^, although this observation has also been disputed^23,24,27^. The uncertainty of whether monomeric Aβ is a viable drug target is caused, in part, by a lack of understanding of the molecular properties of monomeric Aβ and how to stabilise this peptide with specific small molecules that have the potential to be developed as drugs.

The complexity of targeting monomeric Aβ is caused, in part, by the fact that Aβ is intrinsically disordered, as it lacks a well-defined structure and instead exists as a heterogeneous ensemble of conformationally distinct states^28^. The dynamic nature of disordered proteins, and the consequent absence of stable and persistent binding pockets, implies that they do not readily lend themselves to conventional mechanisms of drug-binding, such as the well-established lock-and-key paradigm, in which a drug fits snugly into a single, well-defined binding site.^29-31^ As a result, targeting disordered proteins with small molecules has not been considered a promising drug discovery strategy, and there are no small molecules on the market directly targeting disordered regions despite their high prevalence in disease^2^. A deeper understanding of the possible mechanisms by which small molecules can modify the behaviour of disordered proteins may open new avenues for drug development, not only against Alzheimer’s disease and other neurodegenerative disorders but also many other medical conditions involving disordered proteins, including type II diabetes, and certain forms of cancer and cardiovascular disease^28,29^. This insight may also be particularly valuable in modulating liquid-liquid phase separation, which often involves protein disorder.^32^

Using experimental and computational biophysical techniques and mathematical modelling, we characterise the interaction of the small molecule 10074-G5 (biphenyl-2-yl-(7-nitro-benzo[1,2,5]oxadiazol-4-yl)-amine, **Figure 1a**), with Aβ42 in its disordered, monomeric state. 10074-G5 has been previously identified to inhibit c-Myc-Max heterodimerization^33^ specifically by binding and stabilizing the intrinsically disordered c-Myc monomer^34,35^. Here, we also observe that 10074-G5 binds monomeric Aβ42, a disordered peptide unrelated to c-Myc. As a result of this interaction, 10074-G5 significantly delays both primary and secondary nucleation pathways in Aβ42 aggregation. We characterise this interaction using biophysical experiments and integrative structural ensemble determination techniques, and observe that Aβ42 remains disordered in the bound form with decreased hydrophobicity. Notably, we also observe that the conformational entropy of Aβ42 increases upon interacting with 10074-G5, suggesting that exploiting this phenomenon may be a potential therapeutic strategy for disordered proteins. We further show that this molecule inhibits the pathogenesis associated with Aβ42 aggregation in a *Caenorhabditis elegans* model of Aβ42-mediated toxicity^36^ in a manner consistent with the binding mechanism described *in silico* and characterised *in vitro*.

**Figure 1.**
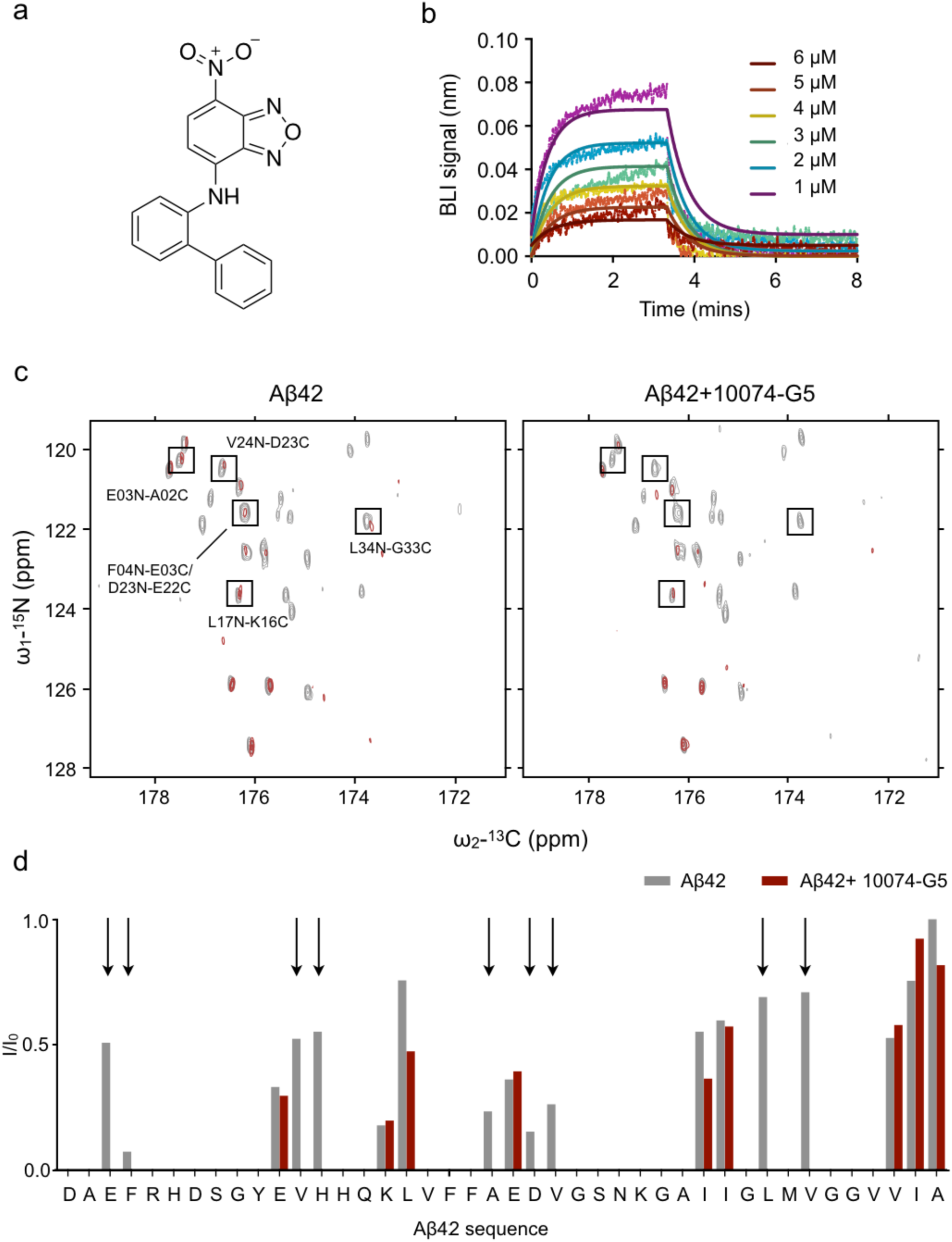
Characterisation of the interaction of 10074-G5 with monomeric Aβ42. **(a)** Structure of biphenyl-2-yl-(7-nitro-benzo[1,2,5]oxadiazol-4-yl)-amine, also known as 10074-G5. **(b)** Biolayer interferometry measurements showing the binding of 10074-G5 to an Aβ42-functionalised surface at various concentrations of the added compound. The curves were corrected for nonspecific binding and baseline drift. Global fitting to simple one-phase association and dissociation equations yields association (*k*_on_) and dissociation (*k*_off_)rates to be 1.5 × 10^3^ M^-1^sec^-1^ and 3.2 ⨯ 10^−2^ sec^-1^, respectively, corresponding to an affinity constant (*K*_D_) of about 21 μM. For this fit, all six curves were constrained to single, shared *k*_on_ and *k*_off_ values. **(c)** 2D H^N–BEST^CON spectra in the absence (left) and presence (right) of 1:2 Aβ42:10074-G5 with (red) and without (grey) selective water pre-saturation, performed at 15 °C. **(d)** Quantification of the relative *I*/*I*_0_ intensities from (c) shows that the peptide amide groups are more exposed to solvent in the presence of 10074-G5. Arrows highlight regions along the sequence in which signals are detectable in the absence of the compound, but not in its presence, thus suggesting that 10074-G5 increases the solvent exposure of specific regions of Aβ42.

## Results

### Selection of the system

We selected the compound 10074-G5 as model system to understand whether and how a small molecule inhibits the aggregation of Aβ by binding the monomeric form of this peptide. We used this molecule as it has been reported to bind the oncogenic disordered protein c-Myc in its monomeric form, and it contains a nitrobenzofurazan moiety, which has been previously shown to inhibit the aggregation of Aβ^37^.

### Characterisation of the binding of 10074-G5 to monomeric Aβ42

We characterised the binding of 10074-G5 with monomeric Aβ42 using a multidisciplinary approach based on experiments and integrative structural ensemble determination. First, we carried out bio-layer interferometry (BLI, see **Materials and Methods**) measurements to characterise this interaction in real-time. We immobilised N-terminally biotinylated monomeric Aβ42 on the surface of super streptavidin sensor tips (**Materials and Methods**) and exposed them to varying concentrations of the small molecule (**Figure 1b**). We observed a concentration-dependent response, indicative of binding. By globally fitting the binding curves to simple one-step association and dissociation equations, such that the fit constrains all curves to share single association (*k*_on_) and dissociation (*k*_off_) rates, we determined *k*_on_ to be 1.5 x 10^3^ ±0.2 x 10^3^ M^-1^sec^-1^ and *k*_off_ to be 3.2 × 10^−2^ ±0.3 × 10^−2^ sec^-1^, corresponding to a binding dissociation constant (*K*_D_) of 21 μM. This affinity value is comparable to other small molecule interactions with disordered proteins^34^.

We then investigated the binding of 10074-G5 and monomeric Aβ42 at the ensemble-averaged, single residue level. To do so, we performed 2D H^N–BEST^CON nuclear magnetic resonance (NMR) experiments^38^ on uniformly ^13^C, ^15^N-labelled monomeric Aβ42 in the presence of 1- and 2-fold concentrations of 10074-G5. As monomeric Aβ42 is relatively stable in solution at low concentrations and temperatures, we examined the binding of 10074-G5 to monomeric Aβ42 under these conditions (20 μM of Aβ42 of at 5 °C). Minimal chemical shift perturbations were observed in the 2D H^N–BEST^CON spectra upon the addition of compound at a 2:1 ligand:protein ratio at 5 °C (**Figure S1**), consistent with other reports of small-molecule binders of disordered proteins^34,39^, suggesting that Aβ42 remains disordered in the presence of 10074-G5. We then performed this experiment at 15 °C in the absence and presence of pre-saturation of the solvent. This experiment, which relies on heteronuclear direct detection with minimal perturbation of proton polarization, provides a valuable tool to study solvent exposed systems in which amide protons experience fast hydrogen exchange^38^. In particular, the signals of amide nitrogen atoms become attenuated when their directly bound protons are in fast exchange with the solvent. After testing this on a well characterized protein (ubiquitin), we performed this experiment on Aβ42. In the absence of 10074-G5, we observed that several hydrophobic residues, particularly C-terminal residues are protected from solvent exchange (Leu17, Leu34, Val36, Ile41, Ala42, see *I*/*I*_0_ values as shown in **Figure 1c,d**). Notably, in the presence of 10074-G5, we observed the quenching of several residues across the sequence of the monomeric Aβ42 peptide that were not quenched in the absence of 10074-G5 (**Figure 1c,d**), suggesting that some residues have increased solvent exposure in the presence of the small molecule. This change in the solvent exchange profile suggests that 10074-G5 interacts with monomeric Aβ42 in a manner that increases the solubility of at least some of the conformations within the monomeric structural ensemble^29,30^.

To obtain further insight into the thermodynamic properties of this interaction, we quantified the heat changes upon 10074-G5 binding to Aβ40 using isothermal titration calorimetry methods (**Figure S2)**. In these experiments, we used Aβ40 instead of Aβ42 because of the higher solubility of Aβ40; we have, however, shown that 10074-G5 has similar effects on the aggregation of Aβ40 as on that of Aβ42 (**Figure S3a)**. The observation of minimal heat changes (**Figure S2)** suggests that the interaction of 10074-G5 with monomeric Aβ is likely to be entropic, as found for the interactions of another small-molecule with a disordered peptide^40^.

To obtain a structural description of how 10074-G5 affects the disordered structural ensemble of Aβ42, we employed metadynamic metainference, an integrative structural ensemble determination approach^41,42^ that combines all-atom molecular dynamics simulations with NMR chemical shift data to improve force field accuracy (see **Materials and Methods, Figures 2 and S4-8**). These simulations reveal that Aβ42 remains disordered in the form bound to 10074-G5, retaining several ensemble-averaged properties. In particular, we noted that most average inter-residue contacts of the unbound peptide remain the same in the bound form (**Figure 2a**). Furthermore, the radii of gyration of the bound and unbound forms of the peptide are also highly similar (**Figure 2b**). The presence of 10074-G5 does, however, alter the conformational ensemble of Aβ42, promoting conformations with lower relative hydrophobic surface area (the fraction of accessible hydrophobic surface area with respect to the total accessible surface area, **Figure 2c**). To determine which residues became more exposed or protected in the presence of 10074-G5, we compared the residue-specific average solvent accessible surface areas per residue in the presence and absence of 10074-G5. Notably, we observe that Lys_34_ and Met_35_ become more protected in the presence of the small molecule. Met_35_ has previously been identified as a key residue for attention of aggregation, as oxidation has been shown to reduce the lag time of primary nucleation^43^.

**Figure 2.**
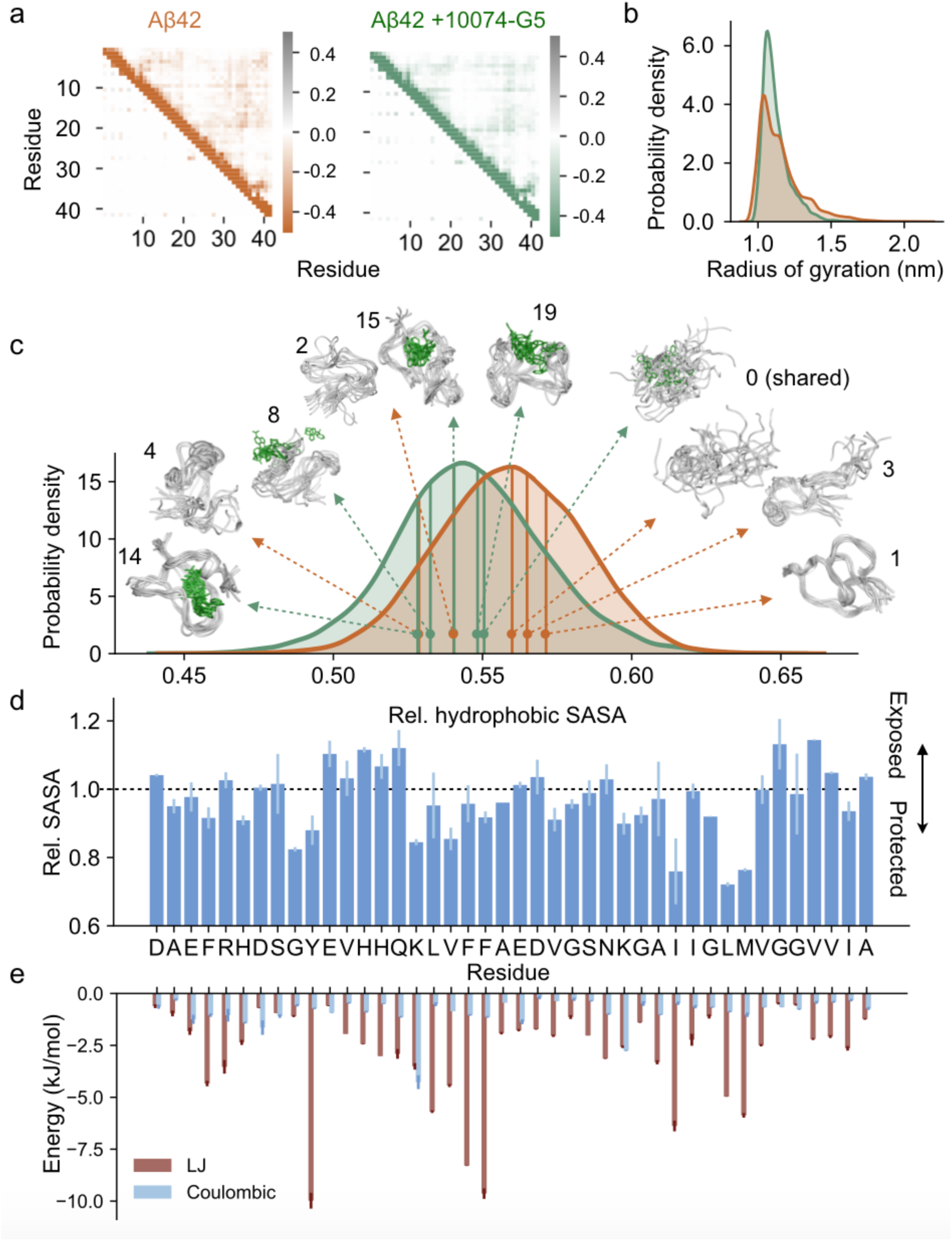
Metadynamic metainference simulations characterise the dynamic binding interaction and show how 10074-G5 promotes Aβ42 conformations with less hydrophobicity. **(a)** Metadynamics metainference simulations demonstrate that inter-residue contact maps for Lennard-Jones (upper right) and Coulomb (lower left) potentials for the unbound (orange) and the bound (green) structural ensembles of Aβ42 with 10074-G5 are highly similar. **(b)** Radii of gyration for the unbound and bound structural ensembles are also highly similar (shown using kernel density estimates of 35,000 points each sampled based on metadynamics weights using a Gaussian kernel). **(c)** Relative hydrophobic solvent accessible surface area (SASA) of Aβ42 (total hydrophobic area over total surface area) of the bound and unbound ensembles, showing that 10074-G5 decreases the relative exposed hydrophobicity of Aβ42. The holo ensemble was calculated only on the protein surface, but in the presence of the compound. Data are shown using kernel density estimates as described in (b). Some of the representative structures of Aβ42 (orange) and 10074-G5 (green) from within these distributions are shown. Numbers indicate cluster IDs shown in **Figure S7. (d)** Ratio (bound/unbound) of the ensemble-averaged, total SASA per residue showing regions of Aβ42 that become more exposed or protected in the presence of 10074-G5. SASAs of the bound ensemble were calculated on the protein, in the presence of 10074-G5. Bar plots represent the entire analysed trajectories, accounting for metadynamics weights. Error bars represent standard deviations between first and second halves of the analysed trajectories. **(e)** Ensemble-averaged, residue-specific Lennard-Jones (LJ) and Coulomb interaction energies show that 10074-G5 has strong interactions with aromatic and charged residues. Error bars represent standard deviations between first and second halves of the analysed trajectories.

The decrease in the hydrophobic surface area is confirmed by the calculation of the average number of hydrogen bonds formed by water molecules within the hydration shell (all waters within 0.4 nm of Aβ42). We found that water molecules form 3.075(1) hydrogen bonds on average in the absence of 10074-G5, a number that increases in its presence to 3.096(1), getting closer to the bulk-like value of 3.448(1) under the conditions investigated here; errors are calculated comparing the first and second halves of the trajectories (**Figure S6a**)^44^. As expected, we observed instead no difference between the average number of hydrogen bonds formed by water molecules in the bulk between the apo and holo simulations (**Figure S6b**). To determine whether or not this binding could be characterised by the release of water molecules upon association, we calculated the average number of water molecules in the hydration shell and show that this value is similar with and without association (**Figure S6c**).

To structurally characterise the unbound and bound conformations of Aβ42 and assess convergence, we performed a clustering analysis (**Figures S7, S8**). We observed that while 10074-G5 binds the extended form in a non-specific manner, all other structural clusters show localisation of the compound within well-defined pockets of Aβ42 for specific conformations (**Figure 2c and S7a**) involving hydrophobic, hydrophilic, charged, and polar residues (**Figure S8**). Interestingly, we also observed that the conformational entropy of Aβ42 is increased in the bound form, exhibiting the so-called “entropic expansion” mechanism which may contribute favourably to the binding free energy^29^ (**Figure S7c**). In stark contrast to the “lock and key” binding mechanism, this observation suggests that the identification of small molecules which increase the conformational entropy of the disordered proteins may be a promising therapeutic strategy.

To probe the energetic contributions to this interaction on an ensemble-averaged, residue-specific level, we analysed Lennard-Jones and Coulomb contributions between 10074-G5 and each residue. We observe strong Lennard-Jones interactions, particularly between aromatic residues (**Figure 2e**) Tyr_10,_ Phe_19_ and Phe _20_ and 10074-G5. The strongest Coulomb interactions occur at charged residues Lys_16_ and Lys_28_ (**Figure 2e**).

### The small molecule 10074-G5 sequesters monomeric Aβ42 and inhibits its aggregation

We measured the kinetics of Aβ42 aggregation at a concentration of 1 μM in the presence and absence of increasing concentrations of 10074-G5. Measurements were performed by means of a fluorescent assay based on the amyloid-specific dye thioflavin-T (ThT), which reports on the overall fibril mass formed during the aggregation process^5,8,16,45-47^. We found that 10074-G5 has a significant effect on Aβ42 aggregation (**Figure 3a,b**). Specifically, the data show that the final value of the ThT fluorescence, which corresponds to the end point of the aggregation reaction, is dependent on the concentration of the compound (**Figure 3a**). The observation of a significant decrease in the final ThT intensity could be due to several non-mutually exclusive possibilities including: 1) interference of the ThT signal by 10074-G5, 2) formation of soluble off-pathway aggregates, 3) sequestration of Aβ42 during the aggregation process^18^.

**Figure 3.**
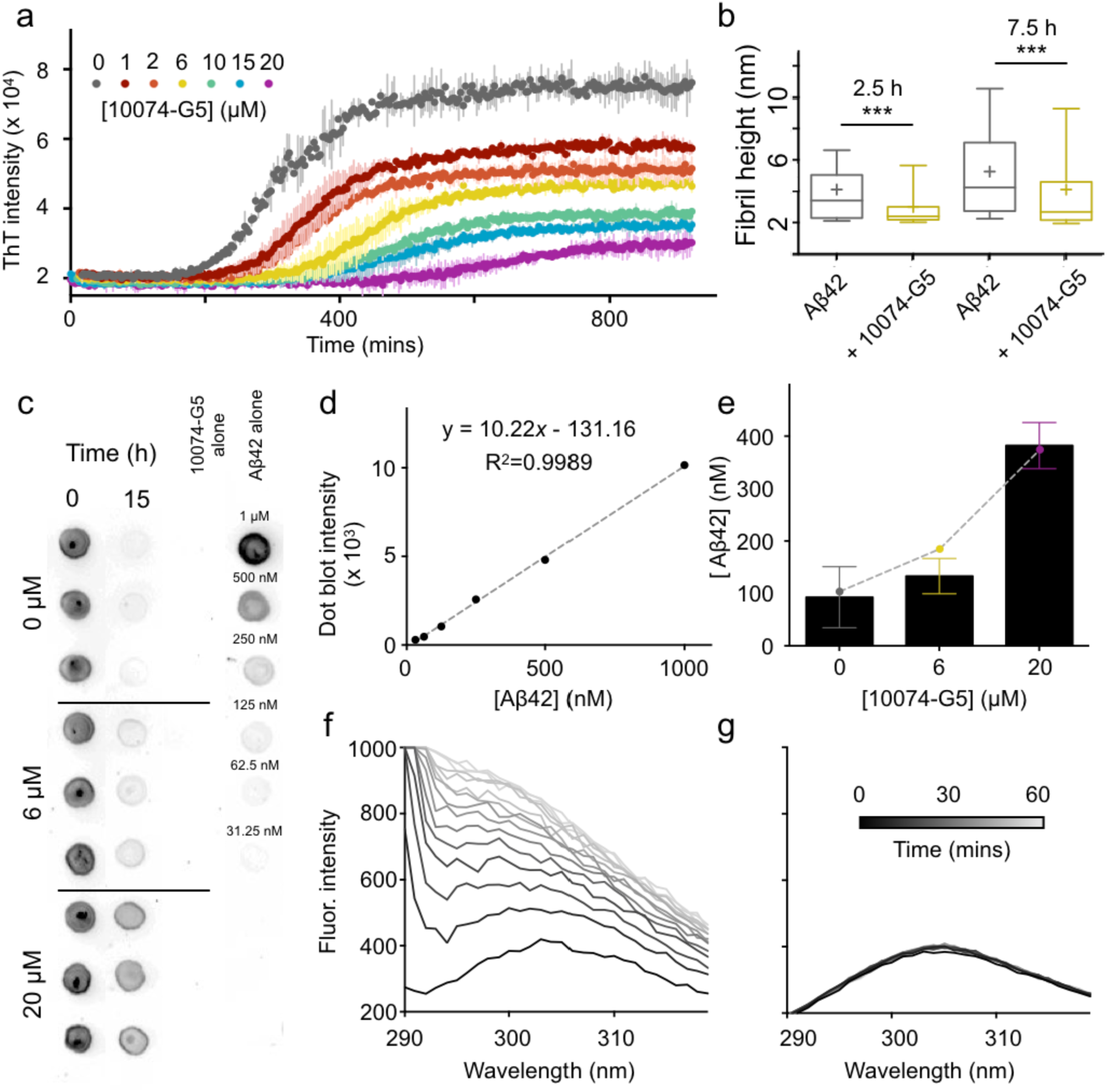
10074-G5 sequesters monomeric Aβ42 and inhibits its aggregation. **(a)** ThT aggregation measurements using 1 μM Aβ42 in the presence of increasing concentrations of 10074-G5 show a concentration-dependent effect of 10074-G5 on Aβ42 aggregation. Error bars represent ±one standard deviation. Measurements were taken in quintuplicate. **(b)** Box-plots of the cross-sectional heights of the Aβ42 fibrils at 2.5 h (N=75 per condition) and 7.5 h (N=200 per condition) formed in the presence and absence of 10074-G5. These data show that 10074-G5 reduces the formation of mature fibrillar aggregates (cross-sectional diameter of ∼6 nm) at given time points (**Figure S9**); boxes indicate median and the standard deviation, a cross indicates the mean, and whiskers show the 10-90 percentile; ***P?<?0.001 by unpaired, two-tailed Student’s t-test. **(c)** Dot blot of soluble Aβ42 before and after the aggregation of 1 μM Aβ42 at 37 °C using the W0-2 antibody in the presence and absence of 10074-G5 indicate sequestration of soluble Aβ42. Blotting was performed in triplicate, as shown. Fit **(d)** and quantification **(e)** used to estimate the concentration of soluble Aβ42 remaining at the end of the aggregation reaction from (d). Error bars represent ±one standard deviation. The dashed line in (e) represents a fit of the dot-blot data to the curve, 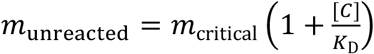 (see Eq. S13), which describes the equilibrium concentration of unreacted monomer from a competitive binding of free monomers to fibril ends and inhibitor molecules (see Materials and Methods). Here, *m*_critical_ = 93 nM is the measured critical concentration of Aβ42, [*C*] is the concentration of 10074-G5 and *K*_D_ = 7 ±1μM is the fitted affinity of 10074-G5 for the soluble material. Intrinsic fluorescence profiles of Tyr10 of 5 μM Aβ42 in the absence **(f)** and presence **(g)** of 1:1 10074-G5 over 1 h show that 10074-G5 delays an increase in fluorescence, suggesting that the compound inhibits early aggregation events including oligomerisation and multimerization of the disordered peptide.

Given the fact that 10074-G5 is a coloured compound, we sought to investigate whether the decrease in the fluorescence intensity of ThT was exclusively due to an interference of 10074-G5 with the dye, or also due to a decrease in the mass of the fibrils formed during the aggregation process. To this end, we performed a ThT-independent dot-blot assay in which we explicitly measured the quantity of soluble Aβ42 over time in the presence and absence of 10074-G5 using the W0-2 antibody, which binds to Aβ (**Figure 3c-e**). The solubility was determined by measuring the amount of Aβ42 that did not sediment after 1 h of ultracentrifugation at 100,000 rpm. We observed that in the presence of a 20-fold excess of 10074-G5, approximately 40% of the total amount of Aβ42 remained in a soluble form (**Figure 3d,e**). These experiments indicate that not all Aβ42 monomers are incorporated in ThT-binding fibrils at the end of the aggregation process, and, thus, that the presence of 10074-G5 sequesters Aβ42 in its soluble form.

These dot-blot data can be explained by an equilibrium model of competitive binding, where monomers can bind both to amyloid fibril ends and to 10074-G5 (**Materials and Methods, Figure 3e**). A fit of the dot-blot data to this equilibrium model (Eq. S13), yields an affinity of 10074-G5 for the monomers of *K*_D_ = 7 ±1 μM, a value broadly consistent with that determined independently from the BLI experiments (*K*_D_ = 21 μM), considering that in the BLI setup Aβ42 is constrained on a surface (**Figure 1c**) whereas the dot blot measurement is performed in bulk solution. We further confirmed the observation that Aβ42 remains soluble by exploiting the intrinsic fluorescence of Tyr10 in the Aβ42 sequence. By monitoring the aggregation of 5 μM Aβ42 from its monomeric form over 1 h, the fluorescence intensity of Tyr_10_ increases considerably (**Figure 3f**) as it becomes buried in a hydrophobic environment in the aggregated state^48^. This experiment reports on early aggregation events that may be invisible to ThT, which is specific for cross-β sheet content as early aggregates such as oligomers or multimers may lack β-sheet structure.^49^ We observed, however that in the presence of an equimolar concentration of 10074-G5, the fluorescence intensity remains constant over time (**Figure 3g**), thereby suggesting that Aβ42 does not self-associate in the presence of 10074-G5.

To further demonstrate that 10074-G5 alters the kinetics of aggregation, we performed 3-D morphological analyses of fibrils using high resolution and phase-controlled^50^ atomic force microscopy (AFM) on the time scale of the aggregation process (**Figures 3b** and **S9**). Single-molecule statistical analysis of the aggregates in the 3-D maps shows that Aβ42, both in the absence and presence of 10074-G5, forms non-mature aggregates with average cross-sectional diameters of approximately 2-3 nm, and mature fibrillar aggregates with average diameters of approximately 5-6 nm, as previously observed^51,52^. It has been previously shown that fibrillar species with diameters less than 6 nm lack a cross-β sheet structure fully stabilised by a tight network of intermolecular hydrogen bonding, as compared to mature fibrillar aggregates^51^. Notably, we observed that at the same time point of aggregation, the fibrillar aggregates formed in the presence of 10074-G5 had smaller cross-sectional diameters than those formed in its absence, with a significantly higher abundance of non-mature species with respect to mature fibrillar species. These data suggest that the process of fibril formation and maturation of cross-β sheet structure in the presence of this compound is considerably slower than in its absence^52,53^ (**Figures 3b** and **S9b**).

### 10047-G5 does not chemically modify Aβ42

To determine whether or not the binding of 10074-G5 to Aβ42 is covalent or induces other chemical modifications, we performed mass spectrometry on Aβ42 incubated in the presence and absence of 10074-G5. Samples were incubated overnight at 37 °C and then spun down using an ultracentrifuge (**Materials and Methods**). The supernatant and resuspended pellet of the aggregation reactions were analysed by matrix assisted laser desorption/ionization (MALDI) mass spectrometry (**Figure S10**). No mass increase was observed following the incubation with 10074-G5, indicating that its presence does not result in detectable covalent chemical modifications to Aβ42.

### 10074-G5 inhibits all microscopic steps of Aβ42 aggregation

In order to better understand the mechanism of inhibition of Aβ42 aggregation by 10074-G5, we performed a kinetic analysis on the ThT aggregation traces. **Figure 4a** shows the ThT kinetic curves normalized relative to the reaction end points. From the normalized data, we observe that 10074-G5 slows down the aggregation reaction in a concentration-dependent manner, consistent with the AFM and dot blot results, confirming a delay in the aggregation process of Aβ42 (**Figure S9**). We then used a chemical kinetics approach^54^ to determine whether the inhibition data could be explained by a monomer sequestration model, in which 10074-G5 inhibits Aβ42 aggregation by binding monomeric Aβ42 and, in this manner, reduces the concentration of monomers available for each microscopic step of aggregation (see **SI**). Specifically, we first fitted the measured aggregation kinetics in the absence of 10074-G5 to a kinetic model of Aβ42 aggregation (see **SI**, Eq. S10)^54^ to estimate the values of the unperturbed rates for primary nucleation, elongation, and secondary nucleation. We then formulated a master equation model for inhibited aggregation kinetics in the presence of 10074-G5 (Eq. S11). We derived explicit integrated rate laws describing inhibited kinetics (Eqs. S11-14 and **Figure S11**), which we used to fit the experimental ThT data in the presence of 10074-G5. For this analysis, we implemented the unperturbed rate constants for aggregation, leaving the value of *K*_D_ as the only fitting parameter. We performed a global fit; all ThT profiles at increasing concentrations of 10074-G5 were not fit individually, but rather using the same choice of *K*_D_, with the dependence on the concentration of 10074-G5 being captured in the integrated rate law through Eq. S14. The result of this global fit is shown in **Figure 4a** and yields an affinity value of *K*_D_ = 40 μM, consistent with other affinities reported for small molecule binders of disordered proteins^34^. It is interesting to note that this estimated affinity is considerably weaker than small molecule binders of structured proteins, which are often in the nM range. It may seem that such a weak binder of Aβ would have little inhibitory effects. However, we note that the level of inhibition in general depends not on the absolute value of *K*_D_, but on the combined parameter *K*_D_/[*C*], where [*C*] is the drug concentration, provided that the rate of binding (*k*_on_[*C*]) is sufficiently fast as compared to the overall timescale of aggregation (*κ*)^55^. Therefore, it is possible to have effective inhibition even for small molecules with *K*_D_ values in the μM range, provided that [*C*] is on the same order of magnitude as *K*_D_ (see **Figure S12**).

**Figure 4.**
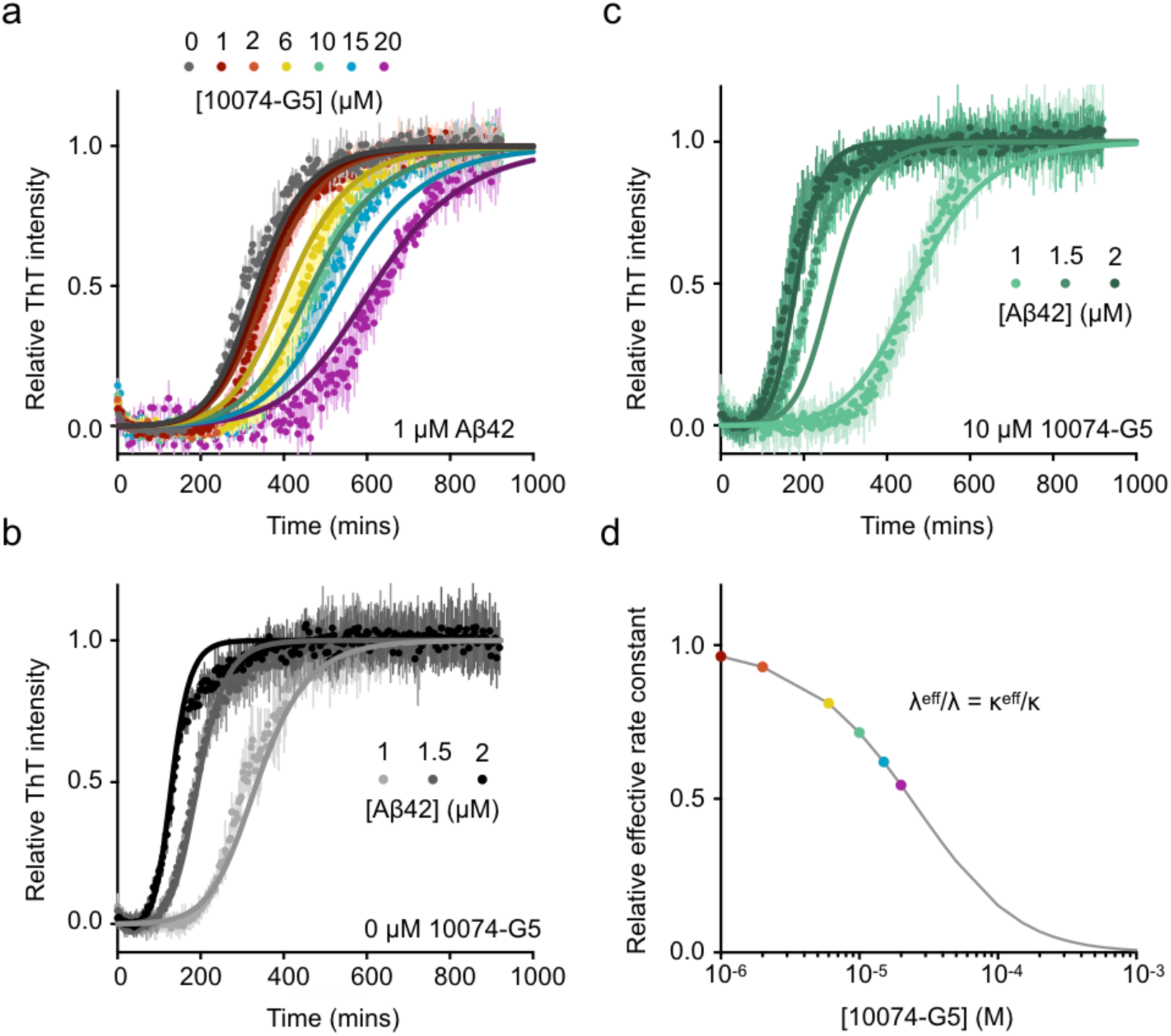
10074-G5 inhibits Aβ42 aggregation primarily by monomer sequestration. **(a)** Global fit of normalized ThT kinetic curves to a monomer sequestration model (Eq. S11), in which 10074-G5 affects the aggregation by binding free monomers. Error bars represent ±one standard deviation. Measurements were taken in quintuplicate. The theoretical curves are obtained using Eq. S10 with unperturbed kinetic obtained from (b) leaving *K*_D_ as the only global fitting parameter (Eq. S14). The global fit yields *K*_D_ = 40 μM. (**b**) Global fit to Eq. S10 of ThT kinetic traces of the aggregation reaction for increasing concentrations of Aβ42 (1, 1.5 and 2 μM) in the absence of 10074-G5. (**c**) Overlay of theoretical kinetic curves from (a) with independent ThT kinetic traces of the aggregation reaction for increasing concentrations of Aβ42 (1, 1.5 and 2 μM) in the presence of 10 μM 10074-G5. Solid curves are predictions of the kinetic monomer sequestration model using the same rate parameters and inhibitor binding constant as in (a) and no fitting parameters. Error bars represent ±one standard deviation. Measurements were taken in triplicate. **(d)** Effective rates of aggregate proliferation through primary (λ) and secondary (κ) nucleation in the presence of varying concentrations of 10074-G5 determined using the global fit in (a).

The analysis of experimental aggregation data in the presence of increasing concentrations of inhibitor using our integrated rate law thus yields an independent method for determining the binding constant of 10074-G5 to the monomers. To provide further support to this analysis, we varied the concentration of monomeric Aβ42 (1, 1.5 and 2 μM) and recorded kinetic traces of aggregation in the absence (**Figure 4b**) and presence of 10 μM 10074-G5 (**Figure 4c**). Using the rate parameters determined from the uninhibited kinetics and the same value of *K*_D_ obtained from the global fit shown in **Figure 4a**, we find that the time course of aggregation predicted by our monomer sequestration model are in good agreement with the independent experimental data (**Figure 4c**).

A key prediction from the monomer sequestration model is that a monomer-interacting compound should interfere with all three microscopic steps of aggregation. In fact, we find that the presence of an inhibitor that binds monomers quickly compared to the overall aggregation rate does not affect the topology of the reaction network. As a result, the inhibited kinetics can be interpreted in terms of effective rates of aggregation that depend on the concentration of inhibitor (Eq. S14). In **Figure 4d**, we show the values of the effective rates of aggregate proliferation through primary (λ) and secondary (κ) nucleation pathways as a function of the concentration of 10074-G5 predicted by this model (see Eq. S10 for a definition of λ and κ). The monomer sequestration model also predicts that the effective rate of elongation should be reduced (**Figure S13b**), although to a lesser extent than the nucleation pathways, which have a stronger monomer concentration dependence. To test this prediction, we performed seeded aggregation experiments in the presence of preformed Aβ42 fibrils to obtain independent measurements of the effective elongation rate as a function of 10074-G5 concentration. We observed that 10074-G5 indeed decreases the effective rate of fibril elongation (**Figure S13a**), consistent with the monomer sequestration mechanism.

We also sought to understand whether 10074-G5 binds monomeric Aβ40 with a comparable affinity to that of Aβ42. To address this question, we applied the monomer sequestration model as used in **Figure 4a**, to fit the inhibitory effects of 10074-G5 on aggregation of 10 μM of Aβ40 (**Figure S3a**). From this analysis, we extracted an affinity constant of 10074-G5 for Aβ40 of 10 μM, a similar value to that obtained for Aβ42 (**Figure S3b**). Given the increased toxicity of Aβ42 as compared to Aβ40, we anticipate the optimisation of small molecules more specific for monomeric Aβ42 over Aβ40.

### Characterisation of the binding of 10074-G5 to stabilised Aβ40 oligomers

Next, we probed whether 10074-G5 alters the behaviour of oligomeric species of Aβ. Although it is extremely challenging to determine whether 10074-G5 modifies the oligomeric intermediates of Aβ42 formed on-pathway to aggregation, which are transient, heterogenous species, it is possible to carry out this analysis more readily on oligomers of Aβ40 stabilised using Zn^2+ 56^. Thus, we next considered whether or not 10074-G5 can alter the behaviour of these stabilised, pre-formed oligomeric species. We incubated pre-formed oligomers in the presence of 10074-G5, centrifuged the samples, and measured the quantities of Aβ40 in the pellet and in the supernatant by using SDS–polyacrylamide gel electrophoresis (SDS-PAGE, **Figure S14a**). The results indicate that these pre-formed oligomers did not dissociate in the presence of 10074-G5. Furthermore, 10074-G5 was found not to alter the turbidity of solutions in which they were present as measured by their absorbance profiles (**Figure S14b**) suggesting that 10074-G5 does not cause such species to change detectably in size. Lastly, dot blots of pre-formed oligomeric samples in the presence and absence of the compound using the OC-antibody, which binds to β-sheets^57^, show that the oligomers maintain their characteristic conformations (**Figure S14c**). Due to the coloured nature of 10074-G5, it was neither possible to characterise the oligomers in the presence of the compound with dynamic light scattering nor analytical ultracentrifugation measurements. Taken together, these data suggest that 10074-G5 does not disaggregate the pre-formed oligomeric species or cause them to undergo further assembly. Nevertheless, it remains possible that this compound affects the evolution of oligomer populations formed during the aggregation reaction, potentially inhibiting their conversion into fibril-competent species.

### 10074-G5 inhibits Aβ42 aggregation in a *C. elegans* model of Alzheimer’s disease

To determine if 10074-G5 can inhibit the formation of Aβ42 aggregates *in vivo*, we tested its effects using a *C. elegans* model of Aβ42-related toxicity (GMC101, **Figures 5, S15, S16**), in which age-progressive paralysis was induced by overexpression of Aβ42 in the body wall muscle cells^36^. The N2 wild-type strain^58^ was used as a control.

**Figure 5.**
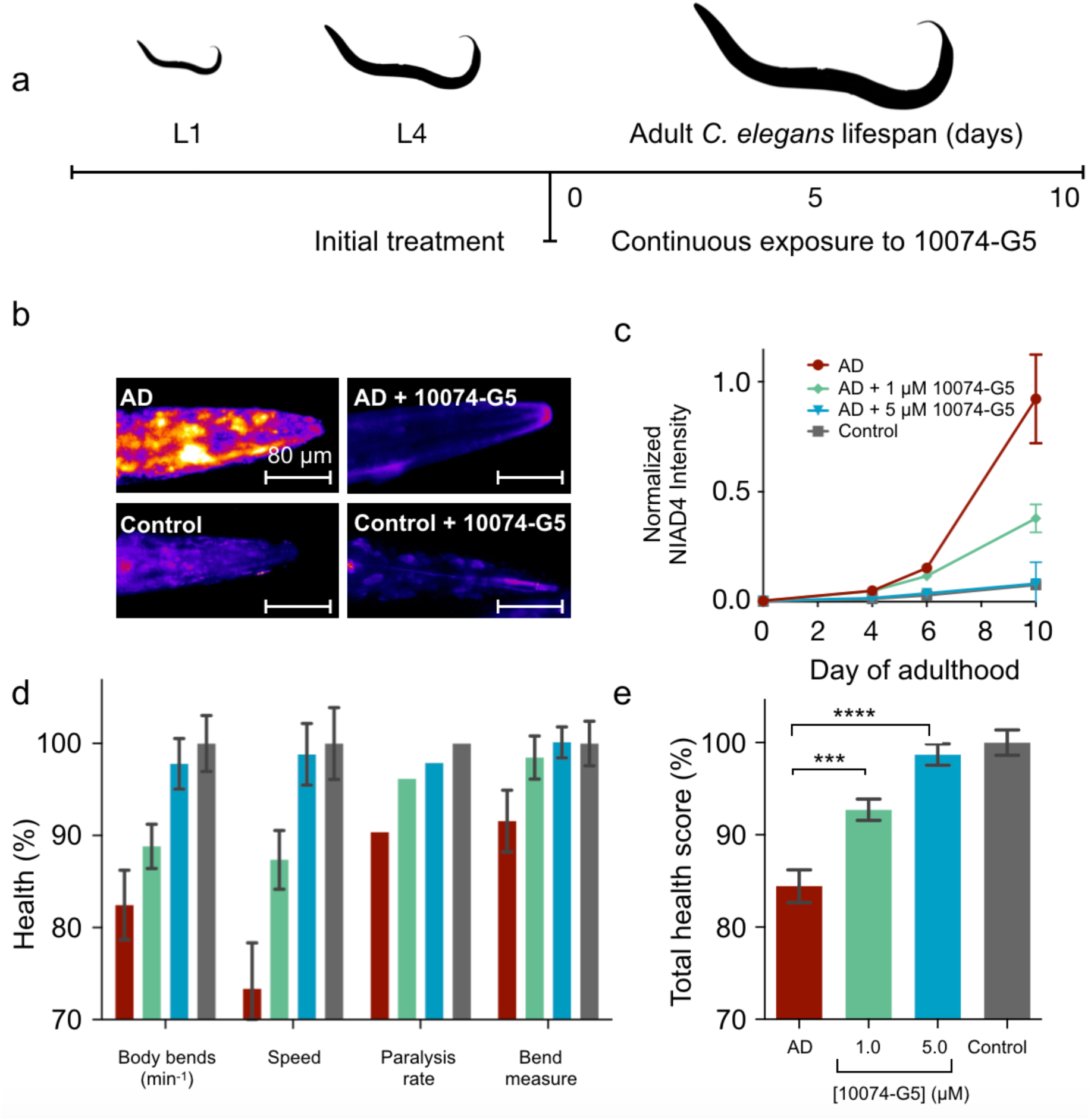
10074-G5 is effective in reducing functional impairment in a *C. elegans* model of Aβ42 toxicity. **(a)** Treatment profile used for the *C. elegans* experiments. **(b)** NIAD-4 staining of *C. elegans* aggregates in the presence and absence of 5 μM 10074-G5. **(c)** Quantification of NIAD-4 intensity shown in panel (b). N=25. **(d)** Health scores (%) for the rate of body bends, speed of movement, paralysis rate, and magnitude of body bends at day 6 of adulthood. The colours are the same as those shown in panel (c). N=150. **(e)** Combined total health scores from panel (d). In all panels, error bars represent ±standard error of the mean (SEM). The symbols *** and **** indicate P < 0.001 and 0.0001, respectively by two-tailed Student’s t-test. 10074-G5 shows minimal movement effects on wild type *C. elegans* (**Figure S16**).

10074-G5 was administered to worms from larval stage L4, and then continuously throughout their lifespan (see **Materials and Methods** and **Figure 5a**). First, we probed the quantity of the aggregates in the animals by means of an amyloid specific fluorescence probe, NIAD-4^20^ (**Figure 5b**,**c)**. The results show that the administration of 10074-G5 resulted in a markedly lower aggregate load. We also monitored a number of phenotypic readouts of worm health, including body bends per minute, the extent of the bending motion, the speed of movement, and also the rate of paralysis. We found that 10074-G5 improved all of these characteristic behavioural parameters in worms expressing Aβ42 in a dose-dependent manner when compared to the untreated worms (**Figure 5d**,**e**).

Taken together, these results demonstrate that the administration of 10074-G5 increases the health of this *C. elegans* model of Aβ42-mediated dysfunction and results in the presence of a smaller number of amyloid aggregates. These findings are consistent with the observation of the inhibition of the aggregation of Aβ42 in the presence of 10074-G5 from the *in vitro* studies (**Figures 3 and 4**). Furthermore, these results suggest that the combination of *in silico, in vitro*, and *in vivo* drug discovery methods holds promise for the identification of novel small molecules which inhibit the toxic behaviour of disease-related disordered proteins.

## Conclusions

We have characterised the binding of the small molecule 10074-G5 to monomeric Aβ42 using a combination of experimental approaches and integrative structural ensemble determination methods. The real-time, dose-dependent responses that we have observed in the BLI experiments demonstrate that 10074-G5 binds Aβ42 in its monomeric form (**Figure 1b**). The NMR experiments and the metadynamic metainference simulations have illustrated that this binding is distinct from most small molecule interactions with structured proteins. In particular, we have observed that 10074-G5 does not bind to a single binding site, but rather binds transiently to many different sites (**Figures 2, S7**). We have also found 10074-G5 induces Aβ42 to adopt many different conformations, keeping it disordered and more solvent-exposed (**Figures 1**) in the bound form. It appears that this interaction is largely entropic (**Figure S2**) and that some of this entropy arises from the increase in conformational entropy of Aβ42 (**Figure S7c**) via the “entropic expansion” mechanism^29^. This structural insight may open new routes for inhibitors of pathogenic disordered proteins and small molecule modulators of liquid-liquid phase separation involving disordered regions.

Additionally, we have characterised the effects of 10074-G5 on amyloid aggregation *in vitro* using a range of biophysical techniques and kinetic theory techniques. This analysis has revealed that as a result of the interaction of 10074-G5 with monomeric Aβ42, this small molecule also reduces the extent to which monomeric Aβ42 contributes to aggregation, thereby effectively slowing down all microscopic aggregation rates. Our kinetic analysis of aggregation inhibition by monomer sequestration highlights that effective inhibition depends on the interplay of two combined parameters, namely a thermodynamic and kinetic one: *K*_D_/[*C*] and *k*_on_[*C*]/*k*_55_. Thus, for a small molecule with given affinity for monomeric Aβ42, increasing *k*_on_ of small molecules for monomeric Aβ42 represents a promising inhibitor optimisation strategy. We also anticipate future work to understand the relationship between *k*_on_ and the change in conformational entropy of Aβ42 towards improved binders of monomeric disordered proteins.

Lastly, we show that 10074-G5 is highly effective at reducing the Aβ42 aggregate load and its associated toxicity in a *C. elegans* model of Alzheimer’s disease.

Collectively, these results indicate the importance of developing a more detailed understanding of the interactions between disordered proteins and small molecules, which in turn could lead to the development of new therapeutic approaches for the many diseases in which such disordered proteins are involved.

## Acknowledgements

GTH is supported by the Gates Cambridge Trust and the Rosalind Franklin Research Fellowship at Newnham College, Cambridge, RL is supported by the Gates Cambridge Trust, FAA by a Senior Research Fellowship Award from the Alzheimer’s Society, UK (Grant Number 317, AS-SF-16-003), TCTM by Peterhouse, Cambridge and the Swiss National Science Foundation, and FSR by Darwin College and the Swiss National Foundation (Grant Numbers P300P2_171219 and P2ELP2_162116, respectively). We acknowledge ARCHER UK National Supercomputing Service under ARCHER Leadership project (Grant Number e510) and PRACE for awarding us access to MareNostrum at Barcelona Supercomputing Center (BSC), Spain for metadynamic metainference simulations. Parameterisation of 10074-G5 was performed using resources provided by the Cambridge Service for Data Driven Discovery (CSD3) operated by the University of Cambridge Research Computing Service (www.csd3.cam.ac.uk), provided by Dell EMC and Intel using Tier-2 funding from the Engineering and Physical Sciences Research Council (capital grant EP/P020259/1), and DiRAC funding from the Science and Technology Facilities Council (www.dirac.ac.uk). MALDI mass spectrometry measurements were performed by Dr. Len Packman at the Protein and Nucleic Acid Chemistry Facility (PNAC) at the Department of Biochemistry, University of Cambridge. The NMR measurements were supported by the iNEXT H2020 Programme (PID: 3017). The work was also supported by the Centre for Misfolding Diseases and the INCEPTION project ANR-16-CONV-0005.

## Data Availability Statement

The data and code that support the findings of this study are available from the corresponding author upon request.

## Materials and Methods

### BLI experiments

A super streptavidin biosensor (ForteBio, Menlo Park, USA) was coated with 15 μg/ml monomeric N-terminally biotinylated Aβ42 (AnaSpec, Fremont, USA) by overnight incubation a solution at 5 °C. Control biosensors were incubated with the same concentration of biocytin. The tips were then rinsed by incubation in buffer for 3 h at room temperature. The binding and dissociation between immobilized Aβ42 and various concentrations of 10074-G5 was monitored for 200 s and 500 s respectively at 37 °C using an Octet Red96 (ForteBio, Menlo Park, USA). The binding of both 10074-G5 to a biocytin-functionalized tip and buffer to a Aβ42-functionlized biosensor were subtracted to account for non-specific binding and baseline drift, respectively. Data were analysed using GraphPad Prism 6. Dissociation data were first globally fit using a one-phase exponential decay to determine the *k*_off_ value. This value was then used as a constraint to determine the global *k*_on_ rate.

*2D H*^*N–BEST*^*CON NMR experiments*. ^13^C, ^15^N uniformly labelled, recombinant Aβ42 peptide (the 42-residue variant lacking the N-terminal M, see ‘*Preparation of recombinant A*β *peptides’*) was purchased from rPeptide and prepared following the manufacturer’s instructions. 20 μM samples were prepared in PBS (pH 7.50), 1% DMSO, with 5% D_2_O (Sigma Aldrich) for the lock. 2D H^N–BEST^CON measurements^38^ were performed at 16.4 T on a Bruker Avance spectrometer operating at 700.06 MHz ^1^H, 176.03 MHz ^13^C and 70.9 MHz ^15^N frequencies, equipped with a triple-resonance cryogenically cooled probehead optimized for ^13^C-direct detection (at the Centro di Risonanze Magnetiche, Florence, Italy). Each 2D H^N–BEST^CON spectrum was acquired with 64 scans. The dimensions of the acquired data were 1024 (^13^C) x 116 (^15^N) points. The spectral width was 29.9 x 33.9 ppm for F_2_ and F_1_, respectively. The relaxation delay was set to 0.3 s. 2D H^N–BEST^CON measurements were repeated with the same parameters except for the inclusion of a weak pre-saturation of the solvent signal during the relaxation delay. Under these conditions, signals of amide nitrogen whose directly bound protons are in fast exchange with the solvent are attenuated. This approach was tested on a well characterized protein (ubiquitin) and then used for the study of the Aβ42 peptide with and without the addition of 10074-G5. 1D ^1^H and 2D BEST TROSY^59^ spectra were acquired before and after measurements were taken to ensure that minimal aggregation had occurred during the course of the measurement. Experimental data were acquired at 5 and 15 °C using Bruker TopSpin 3.1 software and processed with Sparky 3.115.

### Metadynamic metainference simulations

To generate the structural ensembles, we employed an integrative approach that incorporates NMR chemical shift data into molecular dynamics simulations. To this end, we used metadynamic metainference, which compensates for the inaccuracies of the force field, accounts for errors in experimental data, and enhances sampling.^41,42^ All-atom metadynamic metainference simulations^41^ of the unbound and bound form of Aβ42 were performed using GROMACS 2016.4^60^ equipped with the open-source, community-developed PLUMED library^61^ version 2.5^62^, the CHARMM22* force field^63^ and TIP3P water model^64^. The initial conformation of Aβ42 was prepared as a linear peptide using PyMol^65^. A preliminary *in vacuo* molecular dynamics simulation was performed for 1 ns to collapse the extended conformation. This structure was solvated in a rhombic dodecahedron box with an initial volume of 362 nm^3^ containing 11746 water molecules. The solvated system was minimised using the steepest descent algorithm with a target maximum force of 1000 kJ mol^-1^ nm^-1^. A pool of 48 initial conformations was extracted from a preliminary 2 ns simulation at 600 K in the NVT ensemble. Equilibration was then performed in the NVT ensemble for 500 ps at 278 K using the Bussi-Donadio-Parrinello thermostat^66^ and for 500 ps at 278 K in the NPT ensemble using Berendsen pressure coupling^67^ with position restraints on heavy atoms. Production runs were executed in the NPT ensemble at 278 K using the Parrinello-Rahman barostat^68^. A time step of 2 fs was used together with LINCS constraints on all bonds^69^. The van der Waals interactions were cut off at 1.2 nm, and the particle-mesh Ewald method was used for electrostatic interactions^70^. Bound simulations were performed as described above, using the starting structures obtained from the NVT equilibration at 600 K. The 10074-G5 molecule was added to a corner of the box and the system re-solvated with 11734 water molecules. The system was then minimized and equilibrated using the procedures described above. Preliminary parameters for 10074-G5 were taken from the CGenFF software^71,72^, and those with any penalty were explicitly re-parameterised using the Force Field Toolkit^73^ and Gaussian 09^74^ (see **SI** and **Figure S4**).

Chemical shifts were back-calculated at each time step using CamShift^75^ (**Figure S5**). Given that the error of the CamShift predictor is greater than the chemical shift perturbations upon addition of the compound, the same chemical shifts were used to restrain both the unbound and bound simulations. A Gaussian noise model with one error parameter per nucleus type was used in the metainference setup, along with an uninformative Jeffreys prior for each error parameter^41^ (see **SI**). The metainference ensembles for the unbound and bound simulations were simulated using 48 replicas each.

Parallel bias metadynamics^76^ with the well-tempered^77^ and multiple-walkers^78^ protocols was performed using a Gaussian deposition stride of 1 ps, an initial height of 1.2 kJ/mol, and bias factors of 24 and 49 for the unbound and bound simulations, respectively. In the unbound simulations, we used 6 collective variables (CVs) to enhance the conformational sampling of Aβ42 (see **SI**). In the bound simulations, we also included 14 CVs to enhance the conformational sampling of contacts between the compound and the peptide, and 4 CVs to enhance sampling of soft dihedrals in the small molecule (see **SI**). Unbound and bound simulations were run for an accumulated time of 27.8 and 28.2 μs, respectively until convergence was reached (see **SI** and **Figure S7b**). For details on the analysis, see **SI**.

### Preparation of recombinant Aβ peptides

Recombinant Aβ(M1-42) (MDAEFRHDSGY EVHHQKLVFF AEDVGSNKGA IIGLMVGGVVIA) and Aβ(M1-40) (MDAEFRHDSGY EVHHQKLVFF AEDVGSNKGA IIGLMVGGVV), here referred to as Aβ42 and Aβ40, respectively, were prepared by expression in *Escherichia coli* BL21 (DE3) Gold Strain (Agilent Technologies, Santa Clara, USA)^46^. The resulting inclusion bodies were dissolved in 8M urea, ion exchanged in batch mode on diethylaminoethyl cellulose resin, lyophilized, and then further purified with a Superdex 75 HR 26/60 column (GE Healthcare, Chicago, USA). Those fractions containing the recombinant protein, as determined by SDS–polyacrylamide gel electrophoresis, were combined and lyophilized again. To ensure we were working with highly purified monomeric species containing extremely low quantities of aggregated forms of the peptides, size exclusion chromatography was performed directly before the experiments were performed. Aβ40 and Aβ42 solutions were prepared by dissolving the lyophilized peptide in 6 M GuHCl and incubating on ice for 3 h. The solutions were then purified using a Superdex 75 Increase 10/300 GL column (GE Healthcare, Chicago, USA) at a flow rate of 0.5 ml/min and eluted in 20 mM sodium phosphate buffer (pH 8) supplemented with 200 μM EDTA. The center of the peak was collected and the concentrations of the peptides were determined from the integration of the absorbance peak using ε_280_=1495 liter mol^-1^ cm^-1^.

### Preparation of small molecules

10074-G5 was obtained from Sigma Aldrich (St. Louis, MO, USA). The molecules were dissolved in 100% DMSO and then diluted in solutions of Aβ40 or Aβ42 to reach a maximum final DMSO concentration of 1.5%. The total DMSO concentration was matched in the control solutions in all experiments.

### ThT aggregation kinetics

Monomeric Aβ40 or Aβ42 were diluted with buffer and 20 μM ThT from a 2 mM stock and increasing amounts of 10074-G5. Samples were prepared using LoBind Eppendorf tubes (Sigma Aldrich, MO, St. Louis, USA) on ice. Fibrils for seeding experiments were prepared by incubating monomeric Aβ42 at 37 °C overnight. The concentration of fibrils (in monomer equivalents) was assumed to be the initial concentration of monomer. These preformed fibrils were added to a freshly prepared monomer solution to give a final concentration of 15% fibrils.

Samples with or without seed fibrils were pipetted into multiple wells of a 96-well half-area, low-binding polyethylene glycol coating plate (Corning 3881, Corning, USA) with a clear bottom, at 90 μl per well. Plates were sealed with aluminium sealing tape (Corning) to prevent evaporation and then placed at 37 °C under quiescent conditions in a plate reader (CLARIOstar; BMG Labtech, Ortenberg, Germany). ThT fluorescence was measured through the bottom of the plate using 440-nm and 480-nm excitation and emission filters, respectively. ThT fluorescence was followed in quintuplicate for each sample. For analysis of ThT kinetics see **SI**.

### Mass spectrometry

Monomeric Aβ42 was diluted in the aggregation buffer (described above) to a concentration of 15 μM in the presence and absence of 30 μM 10074-G5. Samples were incubated overnight at 37 °C under quiescent conditions to mimic the aggregation experiments. The samples were then spun down using an ultracentrifuge at 100,000 rpm for 1 h at 25 °C to separate the supernatant and pellet. 6 M GuHCl was used to dilute the supernatant by 50% with and resuspend the pellet. Samples were analysed by MALDI mass spectrometry at the Protein and Nucleic Acid Chemistry Facility (PNAC) at the Department of Biochemistry, University of Cambridge.

### Dot-blot assay

Blotting was performed using the Aβ42 sequence-specific antibody (W0-2, MABN10, Millipore, Burlington, USA). Samples were removed from a solution containing 2 μM Aβ42 in the presence and absence of three- and ten-fold equivalence of 10074-G5. To ensure only the monomer was placed on the blots, samples were spun down using an ultracentrifuge at 100,000 rpm for 1 h at 25 °C using a TLA100 rotor. 2 μL of the supernatant were pipetted onto a nitrocellulose membrane (0.2 μM; Whatman). After drying, the membrane was blocked with 5% (w/v) bovine serum albumin (BSA) in phosphate-buffered saline (8 mM Na_2_HPO_4_, 15 mM KH_2_PO_4_, 137 mM NaCl, 3 mM KCl, pH 7.4, PBS) overnight at 5 °C, followed by three 15 min washes with PBS at room temperature. The membrane was then immunised with a 1/1000 dilution of WO-2 anti-Aβ antibody in PBS with 5% BSA overnight at 5 °C, followed by three 15 min washes with PBS at room temperature. The membrane was then incubated for 2 h at room temperature in PBS supplemented with 0.05% tween and an anti-mouse-Alexa Fluor 594 secondary antibody conjugate (ThermoFisher Scientific, Waltham, USA) at room temperature, and then washed three times with PBS supplemented with 0.05% tween. Fluorescence detection was performed using Typhoon Trio Imager (GE Healthcare, Chicago, IL, USA). Blots were quantified using ImageJ. Data were fit to a competitive binding equilibrium model between free monomers and fibrils (Eq. S13). In this model, monomers are either free, aggregated (i.e. part of a fibril), or bound to 10074-G5; the binding of the compound to the monomer is described by a single binding free energy. The binding of monomers to fibril ends is stronger compared to the binding of monomers to the inhibitor. The concentration of free monomer in equilibrium with amyloid fibrils (critical concentration) measured in our experiments is *m*_critical_= 93 nM, consistent with other reports.^79^ The equilibrium concentration of unreacted soluble monomer after ultracentrifugation measured at varying inhibitor concentration is fit to Eq. S13 with *K*_D_ as a fitting parameter. This procedure yields *K*_D_ = 7 ±1 μM, as shown in **Figure 3e**.

### Atomic force microscopy

Solutions of 1 μM Aβ42 in the presence and absence of 6μM 10074-G5 were deposited on mica positively functionalized with (3-aminopropyl) triethoxysilane (APTES, Sigma Aldrich, St. Louis, USA) in the absence of ThT. The incubation times were selected based on the results of the chemical kinetics experiments. The mica substrate was positively functionalized by incubation with a 10 μl drop of 0.05% (v/v) APTES in Milli-Q water for 1 min at ambient temperature, rinsed with Milli-Q water and then dried by the passage of a gentle flow of gaseous nitrogen^51^. AFM sample preparation was carried out at room temperature by deposition of a 10 μL drop of protein solution deposited for 2 min to a surface treated with APTES. The samples were rinsed with Milli-Q water, dried with nitrogen gas, and stored in a sealed container until imaging. AFM maps were acquired by means of a NX10 (Park Systems, Suwon, Korea) and a nanowizard2 (JPK Instruments, Berlin, Germany) system operating in tapping mode and equipped with a silicon tip (PPP-NCHR and μmasch) with a nominal radius of 10 nm. Image flattening and single aggregate statistical analysis were performed by SPIP 6 (Image Metrology, Hørsholm, Denmark) software.

### ITC experiments

Isothermal titration calorimetry (ITC). Measurements were performed using an MicroCal Auto-ITC 200 (GE Healthcare, Chicago, USA) at 15°C. Due to the poor solubility of 10074-G5, monomeric Aβ40 (200 μM) was injected 10 times into a solution containing 7 μM of 10074-G5. All solutions were prepared in phosphate buffer (described above) and contained a minimal amount of dimethyl sulfoxide (DMSO, 0.2%) to ensure that the compound was soluble. Each injection was 3.5 μL in volume and was made on 3 min intervals. Heats of dilution, obtained by separately injecting the peptide into buffer and buffer into the solution containing 10074-G5, were subtracted from the final data. The corrected heats were divided by the number of moles injected and analyzed using Origin 7.0 software (OriginLab, Northampton, USA).

### Characterization of the interaction of 10074-G5 with stabilized oligomers

Stabilised oligomers were formed from Aβ40 as previously described^56^. Briefly, 1 mg of lyophilized Aβ40 was dissolved in 300 μL of hexafluoroisopropanol and incubated overnight at 4 °C. After solvent evaporation under nitrogen gas, Aβ40 was resuspended in DMSO to a concentration of 2.2 mM and sonicated twice for 10 min at room temperature. The protein sample was diluted to a final concentration of 100 μM in 20 mM sodium phosphate buffer with 200 μM ZnCl_2_ at pH 6.9. After incubation for 20 h at 20 °C, the solution was centrifuged for 15 min at 15?000 *g* at room temperature. The pellet containing the oligomers was resuspended in 20 mM phosphate buffer at pH 6.9, with 200 μM ZnCl_2_.

Samples containing 20 μM and 10 μM pre-formed Zn^2+^-stabilised Aβ40 oligomers were incubated in the presence and absence of 20 μM 10074-G5 for 1 h. The turbidimetries of the samples were analysed using a plate reader (BMG Labtech, Aylesbury, UK) at 600 nm. Measurements were background subtracted against buffer alone in the absence and presence of compound. The protein content within samples was quantified using the sequence-specific WO-2 antibody (see *Dot-blot assay*). Similarly, the conformations of the oligomers in the presence and the absence of the compound was probed using the conformation-specific OC antibody^57^ (AB2286, Millipore, Burlington, USA) using the protocols described above (see *Dot-blot assay*).

To determine if the oligomers had dissociated after the incubation in the presence of the compound, the samples were centrifuged for 15 min at 15?000 *g*. The pellet was resuspended in 15 μL of 20 mM phosphate buffer at pH 6.9 with 200 μM ZnCl_2_ and analysed along with the supernatant by SDS–polyacrylamide gel electrophoresis.

### C. elegans experiments

The following *C. elegans* strains were used: The temperature-sensitive human Aβ-expressing strain dvIs100 [unc-54p:: A-beta-1–42::unc-54 3′-UTR + mtl-2p::GFP] (GMC101), where mtl-2p::GFP causes intestinal GFP expression and unc-54p::Aβ1–42 expresses the human full-length Aβ42 peptide in the muscle cells of the body wall. Raising the temperature above 20 °C at the L4 or adult stage causes paralysis due to Aβ42 aggregation in body wall muscle^36^. The N2 wild-type strain was used as a control^36,58^.

Standard conditions were used for the propagation of *C. elegans*^*36*^; the animals were synchronized by hypochlorite bleaching, hatched overnight in M9 (3 g/l KH_2_PO_4_, 6 g/l Na_2_HPO_4_, 5 g/l NaCl, 1M MgSO_4_) buffer, and subsequently cultured at 20 °C on nematode growth medium (NGM) (CaCl_2_ 1mM, MgSO_4_ 1mM, cholesterol 5 μg/ml, 250M KH_2_PO_4_ pH 6, Agar 17 g/L, NaCl 3g/l, casein 7.5g/l) plates seeded with the *E. coli* strain OP50. Saturated cultures of OP50 were grown by inoculating 50 mL of LB medium (tryptone 10g/l, NaCl 10g/l, yeast extract 5g/l) with OP50 and incubating the culture for 16 h at 37 °C. NGM plates were seeded with bacteria by adding 350 μl of saturated OP50 to each plate and leaving the plates at 20 °C for 2-3 days. On day 3 after synchronization, the animals were placed on NGM plates containing 5-fluoro-2’deoxy-uridine (FUDR) (75 μM, unless stated otherwise) to inhibit the growth of offspring.

Aliquots of NGM media containing FUDR (75 μM) were autoclaved, poured, seeded with 350 μL OP50 culture, and grown overnight. After incubating for up to 3 days at room temperature, 2.2 mL aliquots of 10074-G5 dissolved in water at different concentrations were spotted atop the NGM plates. The plates were then placed in a sterile laminar flow hood at room temperature to dry. For the final experiments, worms were transferred onto the 10074-G5-seeded plates directly at larval stage L4 and they were exposed to 10075-G5 for the whole duration of the experiment.

To ensure that the presence of 10074-G5 did not affect the OP50 *E. coli* consumed by the *C. elegans*, we performed a growth assay of the *E. coli* directly from the NGM plates in the presence of 10074-G5 or DMSO after one day of incubation at 24 °C (**Figure S15**). *E. coli* from the NGM plates were added to 4 mL of LB media and diluted to an optical density of 0.25. Then, 3 mL of this starter culture was added to 40 mL of sterile LB media, which was incubated at 37 °C and shaking at 180 RPM. Optical density measurements were collected every 30 minutes and the experiment was performed in duplicate.

All *C. elegans* populations were cultured at 20 °C and developmentally synchronized from a 4 h egg-lay. At 64-72 h post egg-lay (time zero) individuals were transferred to FUDR plates, cultured at 24°C to stimulate aggregation, and body movements were assessed over the times indicated. At different ages, the animals were washed off the agar plates with M9 buffer and spread over an OP50 un-seeded 9 cm plate. The swimming worms were visualized by using a high-performance imaging lens and a machine vision camera, after which their movements were recorded at a high number of frames per second (fps) for 30 s or 1 min^80,81^. Body bends were then quantified using a tracking algorithm^81,82^. Briefly, after an initial background subtraction, a second (nonadaptive) thresholding procedure was performed and worms were identified and labelled. The eccentricity, a measure of the ratio of the minor and major ellipse axes, of each tracked worm was then used to estimate the worm bending as a function of time^81,82^. The total health was calculated by summing the mobility, speed, bend measure, and viability of the worms^81,82^. Total health values were normalized using the values of the control worms. At least 150 animals were examined per condition, unless stated otherwise. All experiments were carried out in triplicate and the data from one representative experiment are shown in **Figure 5**. Control experiments to test the effects of 10074-G5 on the movement of N2 wild type *C. elegans* are shown in **Figure S16**. Two-tailed Student’s t tests (unpaired) were used to calculate *P* values. Statistical analysis was performed using the GraphPad Prism 6 software.

To stain the aggregates within the *C. elegans*, live transgenic animals were incubated with 1 μM NIAD-4 (0.1% DMSO in M9 buffer) for 4 h at room temperature^20^. After staining, animals were allowed to recover on NGM plates for about 24 h to allow destaining via normal metabolism. Stained animals were mounted on 2% agarose pads containing 40 mM NaN_3_ as an anesthetic on glass microscope slides for imaging. Images were captured with a Zeiss Axio Observer D1 fluorescence microscope (Carl Zeiss Microscopy GmbH, Jena, Germany) with a 20 × objective and a 49004 ET-CY3/TRITC filter (Chroma Technology Corp, Bellows Falls, USA). Fluorescence intensity was calculated using ImageJ software (National Institutes of Health) and then normalized as the corrected total fluorescence^20,83^. Only the head region was considered because of the high background signal in the intestinal regions. At least 25 animals were examined per condition, unless stated otherwise. All experiments were carried out in triplicate and the data from one representative experiment are shown in **Figure 5**. Two-tailed Student’s t tests (unpaired) were used to calculate *P* values. Statistical analysis was performed using the GraphPad Prism 6 software.

