## Supplemental Information for "Small molecule sequestration of amyloid-β as a drug discovery strategy for Alzheimer’s disease"

*Details of 10074-G5 parameterisation.* Initial parameters for 10074-G5 compatible with the CHARMM22* force field were determined using the CHARMM General Force Field (CGenFF)[1,2]. Parameters with penalties were re-parametrized using the Force Field Toolkit[3] and Gaussian09[4] as follows: Geometry optimisation was performed at the quantum mechanical (MP2/6-31G*) level. Partial atomic charges were derived to reproduce interaction energies and distances with TIP3P water molecules, in a manner consistent with the CHARMM force field. Briefly, to obtain target data, water molecules were positioned either as a hydrogen bond donor or acceptor as previously described[3] and their optimal distances and orientations with 10074-G5 were calculated at the HF/6-31G* level of theory. Then, the partial atomic charges of 10074-G5 were optimised so that the molecular mechanics interaction energy and distances match the quantum mechanical data. Target data in form of the Hessian matrix for bonds and angles were obtained at the MP2/6-31G* level of theory and were used to fit the corresponding bonded molecular mechanics parameters. The dihedral target data were generated by performing relaxed scans of all dihedral angles with CGenFF penalties using 15° intervals at the MP2/6-31G* level of theory. Given the coupled nature of dihedrals in this multi-ring system, dihedral parameters were simultaneously fit to this torsional potential energy surface as previously described[3]. Dihedral fits to target data are shown in **Figure S4**.

*Details of the metainference approach.* Metainference is a Bayesian integrative approach that enables incorporating experimental data into molecular dynamics simulations[5]. The relative weight of experimental data with respect to the molecular dynamics force field is determined by the accuracy, or level of error, in the data. To account for random and systematic errors in the chemical shift data and in the CamShift[6] predictor, we used a Gaussian model of noise

$$\mathrm{p}\left(\mathrm{d}_{n,m}\middle|\{X\},\sigma_{\mathrm{m}}\right) = \frac{1}{\sqrt{2\pi}\sigma_{\mathrm{m}}}\exp\left[-\frac{\left(\mathrm{d}_{n,m}-\bar{\mathrm{d}}_{n,m}(\{X\})\right)^2}{2\sigma_{\mathrm{m}}^2}\right] \quad \text{(S1)}$$

where $\mathrm{d}_{n,m}$ is the $n$-th data point of the $m$-th dataset, $\{X\}$ is the set of the conformations of all replicas, $\bar{\mathrm{d}}_{n,m}(\{X\})$ is the prediction averaged over our metainference ensemble of 48 replicas, and $\sigma_{\mathrm{m}}$ is an uncertainty parameter. Each experimental dataset contains the chemical shifts of a different nucleus. For both the unbound and bound simulations we used 6 datasets for a total of 230 data points (40 $^{13}$Cα chemical shifts, 34 $^{13}$Cβ chemical shifts, 40 $^{13}$C'chemical shifts, 41 $^{1}$Hα chemical shifts, 41$^{1}$HN chemical shifts, and 34 $^{15}$N chemical shifts).

The uncertainty parameter

$$\sigma_{\mathrm{m}} = \sqrt{(\sigma_{\mathrm{m}}^{\mathrm{B}})^2 + \left(\sigma_{\mathrm{m}}^{\mathrm{SEM}}\right)^2} \tag{S2}$$

quantifies the deviation between predicted and measured data and accounts for random and systematic errors as well as errors in the predictor ($\sigma_{\mathrm{m}}^{\mathrm{B}}$), and the statistical error in calculating ensemble averages over a finite number of replicas ($\sigma_{\mathrm{m}}^{\mathrm{SEM}}$). $\sigma_{\mathrm{m}}^{\mathrm{SEM}}$ was initialised to 0.5 and updated with a windowed average calculation of 500 steps[7]. $\sigma_{\mathrm{m}}^{\mathrm{B}}$ was set to an initial value of 9.0 and sampled using a Monte Carlo algorithm in the range from 0.00001 to 10.0, with a maximal trial move equal to 0.1. For each uncertainty parameter, we used an uninformative Jeffreys prior: $\mathrm{p}(\sigma_{\mathrm{m}}) = 1/\sigma_{\mathrm{m}}$.

*Details of the metadynamics setup.* Parallel bias metadynamics[8] was used to enhance sampling of the conformational landscape of Aβ42, in combination with the multiple-walkers protocol[9]. The collective variables (CVs) used were:

1) The total α-helical content as quantified using the ALPHARMSD keyword in PLUMED. This CV is computed by first generating the set of all possible regions of six consecutive residues within the system and then calculating the root-mean-square distance ($\mathrm{RMSD_i}$) between each segment and an idealized α-helical structure. The CV is then calculated using the following switching function to make it differentiable

$$\mathrm{s}_{\alpha\mathrm{Hel}} = \sum_i \frac{1-\left(\frac{\mathrm{RMSD}_i}{\mathrm{RMSD}_0}\right)^8}{1-\left(\frac{\mathrm{RMSD}_i}{\mathrm{RMSD}_0}\right)^{12}} \tag{S3}$$

where the sum runs over all possible α-helical segments and $\mathrm{RMSD}_0 = 0.08$ nm. A Gaussian width equal to 0.64 was used for this CV.

2) The sum of the total parallel and anti-parallel β-sheet content. Total parallel β-sheet content is calculated using the PARABETARMSD keyword in PLUMED. This value is computed by first generating the set of all possible six residue groupings within the system that can form a parallel β-sheet. Within a given protein chain, two segments containing three continuous residues can form a parallel β-sheet if they are separated by a minimum of 3 residues to accommodate a turn. Then, a root-mean-square distance ($\mathrm{RMSD}_i$) between each grouping and an idealized parallel β-sheet is calculated. The total parallel β-sheet content is then calculated using Eq. S3 above, where the sum runs over all potential parallel β-sheets. Similarly, the total anti-parallel β-sheet content is calculated using the ANTIBETARMSD keyword in PLUMED.

This value is computed by first generating the set of all possible six residue groupings within the system that can form an anti-parallel β-sheet. Within a given protein chain, two segments containing three consecutive residues can form an anti-parallel β-sheet if they are separated by a minimum of 2 residues accommodate a turn. Then, a root-mean-square distance ($\mathrm{RMSD}_i$) between each grouping and an idealized anti-parallel β-sheet is calculated. The total anti-parallel β-sheet content is then calculated using the equivalent of Eq. S3 above, where the sum runs over all potential anti-parallel β-sheets. Finally, the CV is computed by summing the total parallel and anti-parallel β-sheet contents. A Gaussian width equal to 0.33 was used for this CV.

3) The radius of gyration calculated on the Cα carbons, using the GYRATION keyword in PLUMED. This CV is defined as

$$s_{\mathrm{Gyr}} = \sqrt{\frac{\sum_{i\epsilon\mathrm{C}_\alpha} m_i |r_i - r_{\mathrm{COM}}|^2}{\sum_{i\epsilon\mathrm{C}_\alpha}^{n} m_i}} \qquad \text{(S4)}$$

where $\mathrm{m_i}$ and $\mathrm{r_i}$ and are the mass and position of the Cα atom of the $i^{\mathrm{th}}$ residue, respectively and $\mathrm{r_{COM}}$ is the coordinate of the center of mass defined as

$$r_{\mathrm{COM}} = \frac{\sum_{i\epsilon\mathrm{C}_\alpha} m_i r_i}{\sum_{i\epsilon\mathrm{C}_\alpha} m_i} \qquad \text{(S5)}$$

A Gaussian width equal to 0.03 nm was used for this CV.

4) The number of contacts between hydrophobic residues. This CV calculates the number of inter-Cβ atom distances in Ala, Cys, Ile, Leu, Met, Phe, Pro, Trp, or Val residues lower than 0.6 nm. It was calculated using the COORDINATION keyword in PLUMED using the following switching function

$$s_{\mathrm{Coord}} = \sum_{i\epsilon A} \sum_{j\epsilon B} \frac{1-\left(\frac{d_{ij}}{\mathrm{d_0}}\right)^6}{1-\left(\frac{d_{ij}}{\mathrm{d_0}}\right)^{12}} \qquad \text{(S6)}$$

Where $d_0 = 0.6$ nm, $d_{ij}$ is the distance between atoms $i$ and $j$, and $A$ and $B$ are the two groups of atoms between which contacts are calculated. For this specific example, $A$ and $B$ are the same group and self-interactions are excluded. A Gaussian width equal to 0.69 was used for this CV.

5) The number of salt-bridges. This CV is calculated as the number of contacts within a 0.6 nm cut-off range between the atoms of $-COO^-$ groups of aspartic or glutamic acids and atoms of the $-NH_3^+$ groups of lysines or the $-C(NH_2)_2^+$ of arginines. This CV is computed using the COORDINATION keyword in PLUMED as defined by Eq. S6 above. A Gaussian width equal to 2.75 was used.

6) The correlation between consecutive $\psi$ torsion angles. This CV was calculated using the DIHCOR keyword in PLUMED using the following equation:

$$s_{\text{Tors}} = \frac{1}{2}\sum_i [1 + \cos(\psi_i - \psi_{i+1})] \quad \text{(S7)}$$

for every residue, i. A Gaussian width equal to 1.34 was used for this CV.

14 CVs were added to the bound simulation to enhance the sampling of contacts between atoms from the 10074-G5 molecule and atoms from consecutive three-residue regions of Aβ42. These were implemented using the COORDINATION keyword in PLUMED using the following equation:

$$s_{\text{Con}} = \sum_{i\epsilon A} \sum_{j\epsilon B} \frac{1-\left(\frac{d_{ij}}{\mathrm{d}_0}\right)^n}{1-\left(\frac{d_{ij}}{\mathrm{d}_0}\right)^m} \quad \text{(S8)}$$

where $d_{ij}$ is the distance between atoms $i$ and $j$, $n = 6$, $m = 12$, and $d_0 = 1.0$ Å. A Gaussian width of 1.0 was used for each of these CVs.

Lastly, 4 CVs were added to enhance the sampling of soft dihedrals of 10074-G5 using the TORSIONS keyword in PLUMED. A Gaussian width of 0.1 was used for each of these CVs. These were applied to the dihedrals in the top four subplots shown in **Figure S4.**

*Details of structural ensemble analysis.* The unbound and bound simulations contain 278,064 and 256,128 frames respectively. Prior to analysis, we checked that no frames in which the peptide interacted with its periodic image, such that the minimum distance between two peptides was less than 1.2 nm, were included. Furthermore, approximately the first 14 μs of the concatenated trajectories were discarded for equilibration. For the bound ensemble, only frames in which 10074-

G5 was in close contact with the peptide were considered, such that the minimum distance between the peptide and 10074-G5 was less than 0.8 nm (253,891 frames).

All observables are calculated as ensemble averages using the weights $w(\mathbf{s}_i)$ obtained from the bias potential $V_{\mathrm{PB}}(\mathbf{s}_i)$ at the end of metadynamics simulations[10] using

$$w(\mathbf{s}_i) = \frac{e^{\frac{V_{\mathrm{PB}}(\mathbf{s}_i)}{k_{\mathrm{B}}T}}}{\sum_j^N e^{\frac{V_{\mathrm{PB}}(\mathbf{s}_j)}{k_{\mathrm{B}}T}}} \qquad \text{(S9)}$$

where $\mathbf{s}_i$ is the value of all CVs at time $i$, $N$ is the total number of time steps, $k_{\mathrm{B}}$ is the Boltzmann constant and $T$ is the temperature.

Contact maps were computed based on Lennard Jones and Coulomb interaction energies between residues calculated using the GROMACS gmx energy tool. Only short-range interaction energies were considered. These interaction energies were then converted to a ternary matrix where in which interactions that were less than one standard deviation from the mean were assigned a value of -1, those within one standard deviation of the mean were assigned a value of 0, and those greater than one standard deviation of the mean were assigned a value of 1.

Hydrophobic surface areas were computed using the GROMACS gmx sasa tool. Hydrophobic atoms were defined as those whose partial charges range from -0.2 to 0.2, all other atoms were defined as hydrophilic. Relative hydrophobic surface areas were computed by normalising over the total solvent exposed surface area.

To characterise the structural similarities and differences between the unbound and bound simulations and to assess their convergence (**Figures S7**), we performed a cluster analysis of the trajectories. Given that the cluster analysis is highly memory-intensive, we sampled a subset of 35,000 frames from each trajectory based on the metadynamics weights. Inter-residue Lennard Jones interaction energies were calculated for each frame and the shuffled data used as input for the GROMOS clustering algorithm[11] using the root-mean-squared-deviation (RMSD) as a measure of similarity between matrices. We used cut-off values equal to 8.0 kJ mol$^{-1}$ to identify clusters and determine their populations. To assess convergence of a given simulation, we calculated the population of the 5 most populated clusters of the unbound and bound ensembles over time **(Figure S7)**. We also calculated the standard deviations of the populations of these states between the first and second halves of the analysed trajectories **(Figure S7b)**. To depict each cluster graphically, 10 frames were randomly selected based on their distance to the cluster centre as defined by the clustering algorithm.

The Lennard-Jones and Coulomb interaction energies between residues and 10074-G5 were computed using the GROMACS gmx energy tool (**Figure 2e**). Only short-range interaction energies were considered. Convergence was analysed by comparing values from the first and second halves of the analysed trajectories.

Hydrogen bonds were calculated using the GROMACS gmx hbond tool. Solvent atoms were defined to be in the shell if they were within 0.4 nm of the protein and/or 10074-G5[12]. All other solvent atoms were considered to be part of the bulk. The number of hydrogen bonds in the shell was computed as the sum of 1) the number of hydrogen atoms between shell and protein atoms ($H_{\mathrm{SP}}$), 2) the number of hydrogen atoms between shell and bulk atoms ($H_{\mathrm{SB}}$), 3) two times the number of hydrogen atoms between all shell atoms ($2 \times H_{\mathrm{SS}}$), and, 4) in the case of the holo simulations, between the shell and small molecule ($H_{\mathrm{SM}}$). This number was then divided by one-third of the total number of atoms in the shell to obtain the average number of hydrogen bonds per water molecule in the shell. Similarly, the number of hydrogen bonds in the bulk was computed was the sum of 1) the number of hydrogen atoms between bulk and shell atoms ($H_{\mathrm{BS}}$), 2) two times the number of hydrogen atoms between all bulk atoms ($2 \times H_{\mathrm{BB}}$), and, 3) in the case of the holo simulations, between the bulk and small molecule ($H_{\mathrm{SB}}$ ). This number was then divided by one-third of the total number of atoms in the bulk to obtain the average number of hydrogen bonds per water molecule in the bulk.

*Kinetic analysis of experimental aggregation data using a monomer sequestration model.* The kinetic traces of Aβ42 aggregation in the absence of the inhibitor are described by the following formula[13]

$$\frac{M(t)}{M(\infty)} = 1 - \left(\frac{B_+ + C_+}{B_+ + C_+ e^{\kappa t}} \frac{B_- + C_+ e^{\kappa t}}{B_- + C_+}\right)^{\frac{k_\infty^2}{\kappa \tilde{k}_\infty}} e^{-k_\infty t} , \quad \text{(S10a)}$$

where $M(t)$ is the concentration of monomer found within the fibrils (proportional to the ThT fluorescence signal), and[13]

$$C_\pm = \frac{\lambda^2}{2\kappa^2} , \quad \text{(S10b)}$$

$$k_\infty = \kappa\sqrt{\frac{2}{n_2(n_2+1)} + \frac{2\lambda^2}{n_c\kappa^2}}\,, \tag{S10c}$$

$$\tilde{k}_\infty = \sqrt{k_\infty^2 - 4C_+C_-\kappa^2}\,, \tag{S10d}$$

$$B_\pm = \frac{k_\infty \pm \tilde{k}_\infty}{2\kappa}\,, \tag{S10e}$$

$$\lambda = \sqrt{2k_+k_\mathrm{n}m(0)^{n_\mathrm{c}}}\,, \tag{S10f}$$

$$\kappa = \sqrt{2k_+k_2m(0)^{n_2+1}}. \tag{S10g}$$

Here, $k_+, k_\mathrm{n}, k_2$ are the rate constants for fibril elongation, primary nucleation and secondary nucleation, respectively. $m(0)$ is the initial monomer mass concentration, and $M(\infty)$ is the fibril mass concentration at the end of the aggregation reaction. $M(\infty)$ is given explicitly by $M(\infty) = m(0) - m_\mathrm{critical}$, where $m_\mathrm{critical} = k_-/k_+$ is the critical monomer concentration and $k_-$ is the rate constant of monomer dissociation from the fibril ends. The parameters $n_\mathrm{c} = 2$ and $n_2 = 2$ are the reaction orders of primary and secondary nucleation[13]. We used Eq. S10a to extract the rate parameters $k_\mathrm{n}k_+$ and $k_2k_+$ from a global fit of independent kinetic traces measured at increasing concentrations of Aβ42 in the absence of the compound (**Figure 4b**). This fit of the unperturbed kinetic curve yields the following values for the combined rate parameters $k_nk_+ = 1.6 \times 10^2 \pm 0.08 \times 10^2\ M^{-2}s^{-2}$ and $k_2k_+ = 5.9 \times 10^{10} \pm 0.1 \times 10^{10} M^{-3}s^{-2}$, consistent with the previous analysis[14].

Next, we developed a monomer sequestration model to describe kinetic traces in the presence of increasing concentrations of the compound (**Figure 4a**). In this model, we hypothesise that a reduction of the concentration of free monomers available for aggregation due to sequestration can account for the observed inhibition data. This situation is captured by the following kinetic equations:

$$\frac{dP(t)}{dt} = k_\mathrm{n}m(t)^{\mathrm{n_c}} + k_2m(t)^{\mathrm{n_2}}M(t)\,, \tag{S11a}$$

$$\frac{dm(t)}{dt} = -(2k_+m(t) - 2k_-)P(t) - k_\mathrm{on}[C]\,m(t) + k_\mathrm{off}m_\mathrm{bound}(t)\,, \tag{S11b}$$

$$\frac{dm_\mathrm{bound}(t)}{dt} = k_\mathrm{on}m(t)[C] - k_\mathrm{off}m_\mathrm{bound}(t)\,, \tag{S11c}$$

where $[C]$ is the concentration of the inhibitor 10074-G5, $m(t)$ and $m_{\mathrm{bound}}(t)$ are, respectively, the concentrations of free and bound monomers; moreover, $k_{\mathrm{on}}$ and $k_{\mathrm{off}}$ are the association and dissociation constants of the inhibitor to/from the monomers, respectively[15]. We work in a regime where the binding and unbinding rates of the inhibitor are fast compared to the overall rate of aggregation $\kappa$. For inhibitor concentrations between 1 μM and 20 μM, the rate of inhibitor binding is $k_{\mathrm{on}}[C] = 1.5 \times 10^{-3} - 3.0 \times 10^{-2}\ \mathrm{s}^{-1}$, while the rate of unbinding is $k_{\mathrm{off}} = 3.2 \times 10^{-2}\ \mathrm{s}^{-1}$. These rates are faster than that of the overall aggregation reaction $\kappa = 3.0 \times 10^{-4}\ \mathrm{s}^{-1}$ (for 1 μM Aβ42), as determined from a global fit of unperturbed aggregation traces to Eq. S10 (**Figure 4b**). Due to this separation of timescales, aggregation in the presence of the inhibitor occurs in two stages (**Figure S11**). First, a rapid pre-equilibrium is established between free and inhibitor-bound monomers, yielding to a fast initial drop in free monomer concentration and a rapid increase in bound monomer fraction (**Figure S11a,b**). The values of free and bound monomer concentrations at the end this rapid pre-equilibrium phase correspond to an equilibrium between free and inhibitor-bound monomers are obtained by setting $\frac{dm_{\mathrm{bound}}(t)}{dt} = 0$ in Eq. S11, yielding

$$\frac{m_{\mathrm{bound}}}{m(0)} = 1 - \frac{1}{1+[C]/K_{\mathrm{D}}} \quad \text{and} \quad \frac{m}{m(0)} = \frac{1}{1+[C]/K_{\mathrm{D}}}, \qquad \text{(S12)}$$

where $K_{\mathrm{D}} = k_{\mathrm{off}}/k_{\mathrm{on}}$ is the affinity of 10074-G5 for the monomer. These expressions correspond to the dashed lines in **Figure S11a,b** and enter as initial conditions for the second, slower phase of aggregation, determining in this manner the extent of inhibition. During the second phase of aggregation, fibrils form from an effectively lower monomer concentration, which is the origin of the inhibitory effect. The aggregation reaction proceeds until equilibrium is reached. The equilibrium concentrations of free and bound monomers are obtained by applying the principle of detailed balance. The equilibrium concentration of free monomers equals the critical monomer concentration, independently of the amount of inhibitor added. The equilibrium concentration of inhibitor-bound monomers is dependent on the inhibitor concentration and is from Eq. S12 as $m_{\mathrm{bound}} = \frac{m_{\mathrm{critical}}[C]}{K_{\mathrm{D}}}$. Summing the critical monomer concentration and the equilibrium concentration of monomers bound to 10074-G5 yields the following expression for the equilibrium concentration of unreacted soluble monomer left at the end of the aggregation reaction:

$$m_{\mathrm{unreacted}} = m_{\mathrm{critical}} + m_{\mathrm{bound}} = m_{\mathrm{critical}}\left(1 + \frac{[C]}{K_{\mathrm{D}}}\right). \qquad \text{(S13)}$$

We can also obtain an explicit integrated rate law describing the time course of aggregation in the presence of the inhibitor (**Figure S11c**). The explicit equation describing the time-varying aggregate mass concentration is found to be identical to Eq. S10, but the parameters $\lambda$ and $\kappa$ are replaced by renormalized ones, given by

$$\frac{\lambda^{\mathrm{eff}}}{\lambda} = \left(\frac{1}{1+[C]/K_{\mathrm{D}}}\right)^{\frac{\mathrm{n_c}+1}{2}} \tag{S14a}$$

and

$$\frac{\kappa^{\mathrm{eff}}}{\kappa} = \left(\frac{1}{1+[C]/K_{\mathrm{D}}}\right)^{\frac{\mathrm{n_2}+1}{2}} . \tag{S14b}$$

Similarly, the effective rate of elongation depends on the compound concentration as:

$$\frac{{k_+}^{\mathrm{eff}}}{k_+} = \frac{1}{1+[C]/K_{\mathrm{D}}} . \tag{S14c}$$

The performance of our integrated rate law against numerical integration of the master equation S11 is shown in **Figure S11c**. We used these explicit expressions to fit globally kinetic traces in the presence of increasing concentrations of the compound. In particular, we used Eq. S10, where the kinetic parameters vary with 10074-G5 concentration according to Eq. S14. This global fit, which has one single fitting parameter $K_{\mathrm{D}}$, yields $K_{\mathrm{D}} = 40$ µM and is shown in **Figure 4a**. To validate these fits, we applied this model to independent kinetic traces measured at increasing concentrations of Aβ42 in the presence of 10µM of the drug (**Figure 4c**). We used Eq. S10 and Eq. S14 with the same parameters as in **Figure 4a** to predict the reduction of monomer concentration available for aggregation due to the presence of 10µM of the drug and predicted the resulting kinetic profiles without introducing any fitting parameters (**Figure 4c**). This analysis was done in Mathematica Version 11.0.0.0 and using the online available Amylofit platform[16].

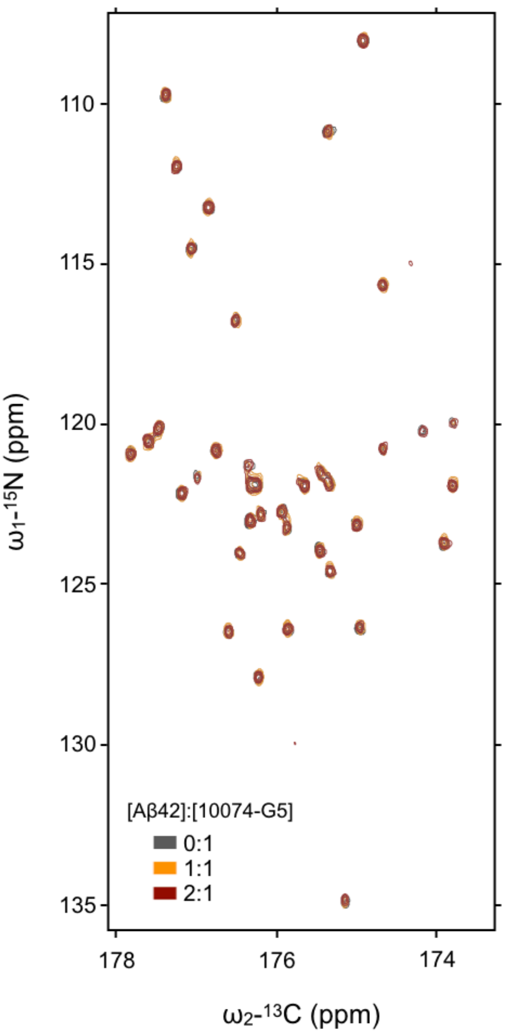


**Figure S1. 2D H$^{N\text{–}BEST}$CON spectra of Aβ42 in the presence of 1- and 2-fold excess 10074-G5**. 2D H$^{N\text{–}BEST}$CON spectra in the absence (grey) and presence of 1- (orange) 2-(red) fold concentration of 10074-G5 at 5 °C.

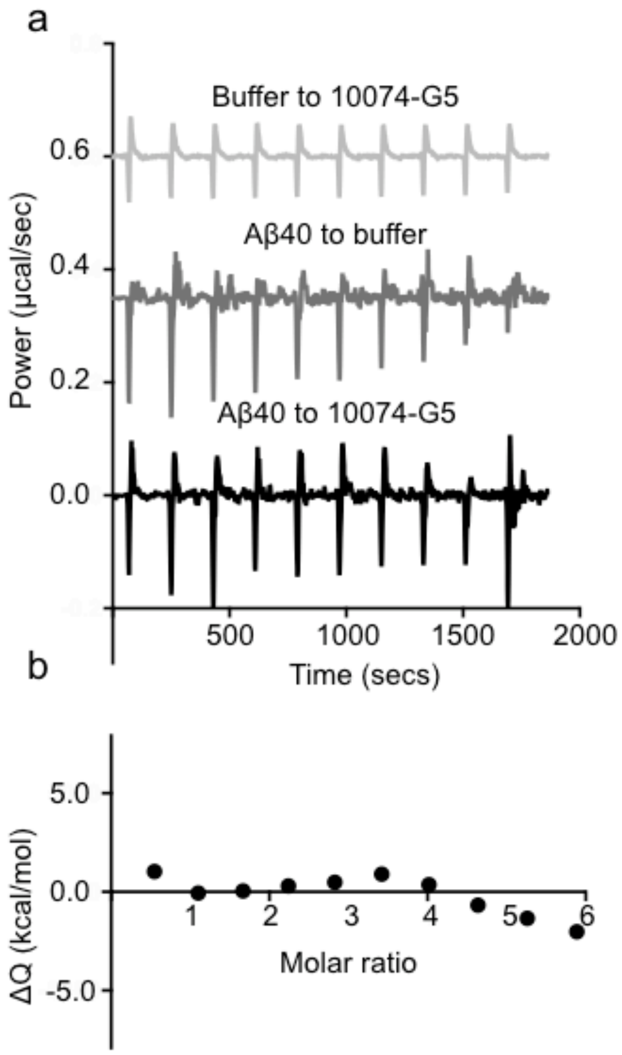


**Figure S2. Isothermal titration calorimetry measurements of the interaction of 10074-G5 with monomeric Aβ40. (a)** Isothermal titration calorimetry experiment of Aβ40 (200 µM) titrated into a solution of 7 µM 10074-G5 with corresponding heats of dilution. **(b)** Integrated peaks from (a) accounting for heats of dilution.

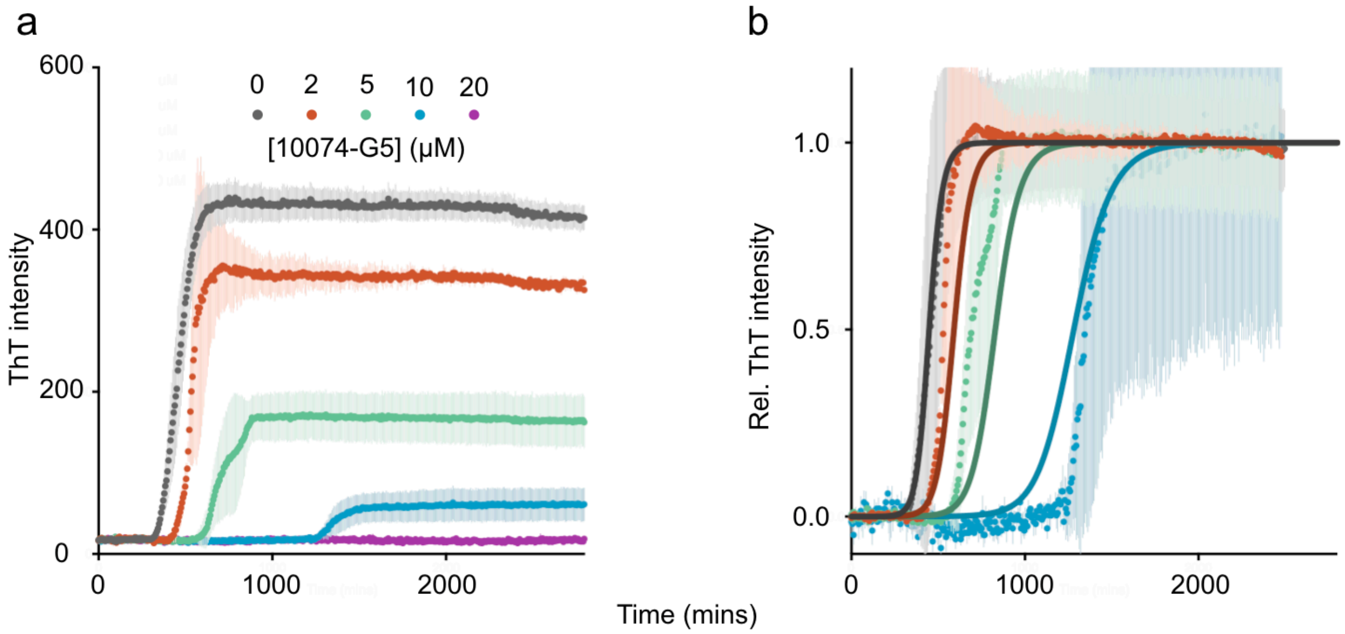


**Figure S3. 10074-G5 inhibits Aβ40 aggregation in an *in vitro* assay. (a)** ThT kinetic traces (see "*ThT aggregation kinetics*" in Materials and Methods) of the aggregation of 10 µM Aβ40 at increasing concentrations of 10074-G5. Error bars represent ± one standard deviation. Measurements were taken in quintuplicate. **(b)** Global fit (solid lines) of normalized ThT kinetic curves (a) to the monomer sequestration model (Eq. S11), in which 10074-G5 affects the aggregation by binding free monomers. The unperturbed rate parameters were obtained by fitting the data without inhibitor to Eq. S10. Kinetic curves in the presence of increasing inhibitor were then globally fit to Eq. S10 with the inhibitor dependence of the perturbed rate parameters given by Eq. S14, leaving $K_D$ as the only global fitting parameter. The global fit yields $K_D = 10$ µM.

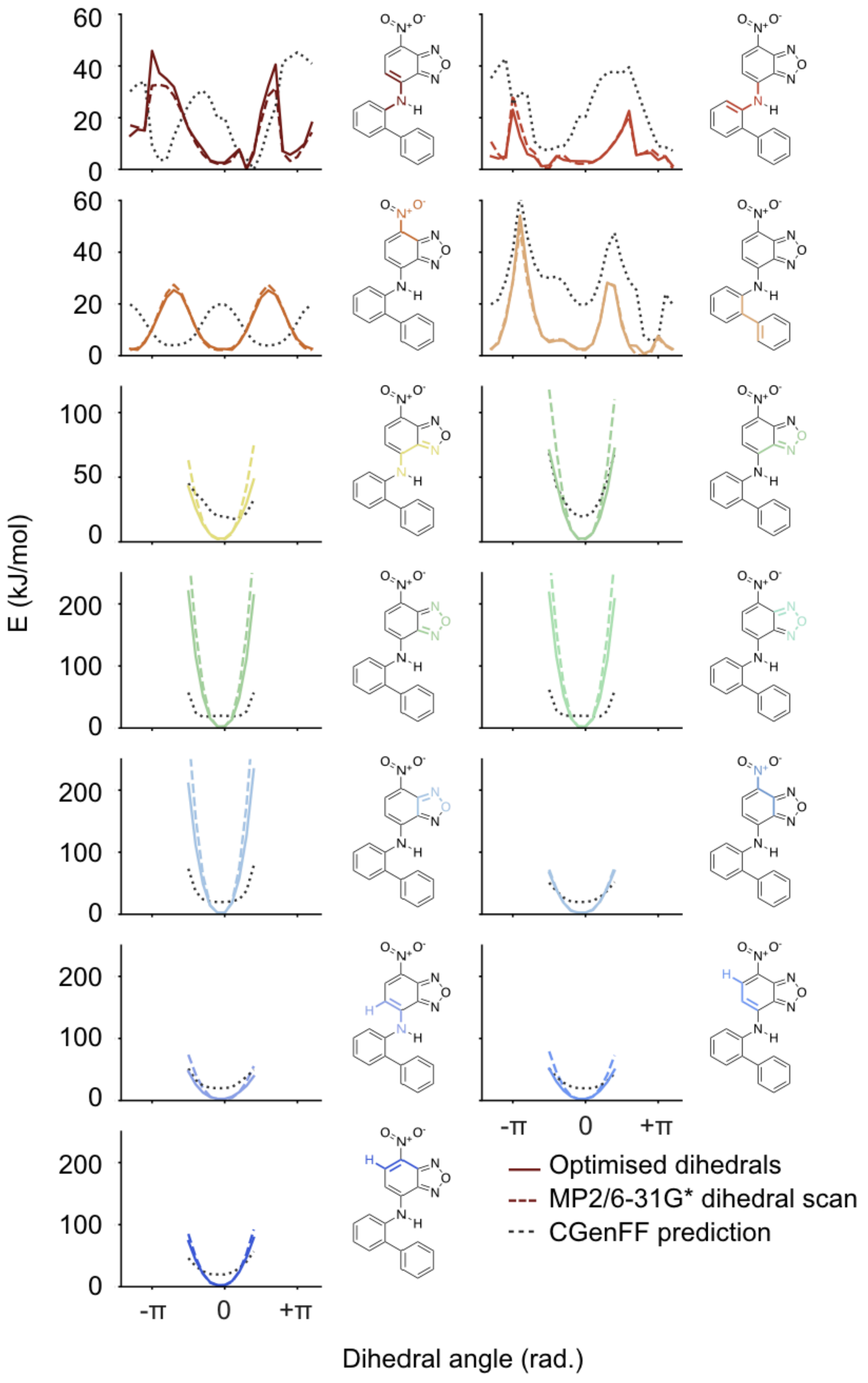


**Figure S4. Parameterisation of the dihedral angles of 10074-G5.** Black dashed lines show the results from the CHARMM general force field (GGenFF)[1,2]. Coloured dashed lines show target data obtained from quantum mechanical dihedral scans performed at the MP2/6-31G* level of theory (see "*Details of 10074-G5 parameterisation*" in SI Materials and Methods). Solid coloured lines show dihedrals after optimisation with the force field toolkit[3]. Coloured atoms and bonds on structures highlight the dihedral of interest for each fit.

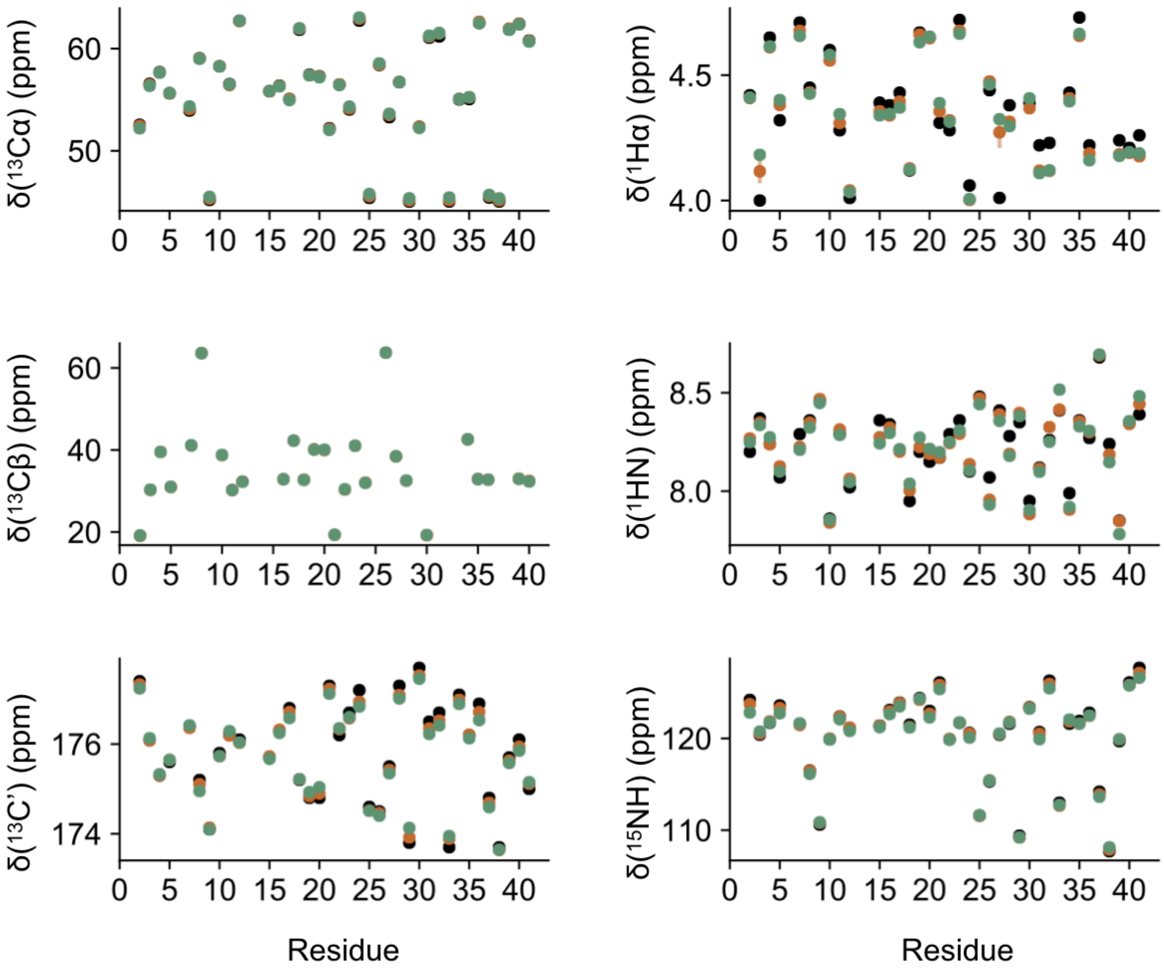


**Figure S5. Consistency of metadynamic metainference simulations with NMR chemical shifts.** Comparison of the NMR chemical shift (δ) data (black) with those predicted from the unbound (orange) and bound (green) simulations. NMR chemical shift data was used to restrain the simulations to improve force field accuracy. Error bars (often smaller than data points) represent standard deviations between the chemical shifts calculated on the first and second halves of the analysed trajectories.

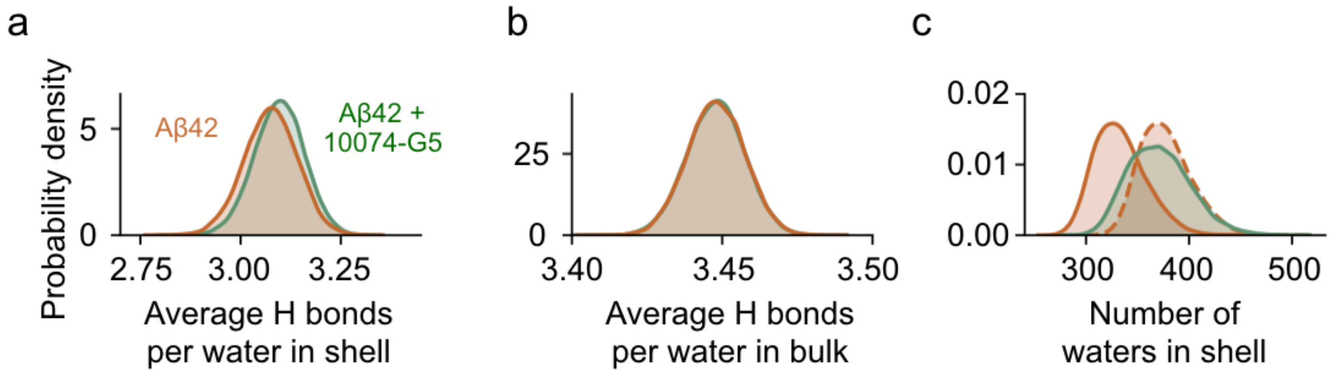


**Figure S6. Characterisation of water properties in metadynamic metainference metainference simulations. (a)** Average number of hydrogen (H) bonds per water molecule in the shell (all water molecules within 0.4 nm of Aβ42 with and without 10074-G5). Distributions are shown using kernel density estimates of 35,000 points each sampled based on metadynamics weights using a Gaussian kernel. The orange curve represents the apo ensemble, the green curve represents the holo ensemble. **(b)** Average number of H bonds per water molecule in the bulk (all water molecules not in the shell) using the same parameters and colors as shown in panel (a). **(c)** Kernel density estimates of the number of water molecule in the shell for the apo (orange curve, solid line) and holo (green curve, solid line) simulations. Using the dissociated frames from the holo simulation (in which 10074-G5 was further than 0.8 nm from the protein), we estimate the number of water molecules in the shell of 10074-G5 alone to be approximately 44.2. The orange dashed distribution represents the apo simulation shifted by this value.

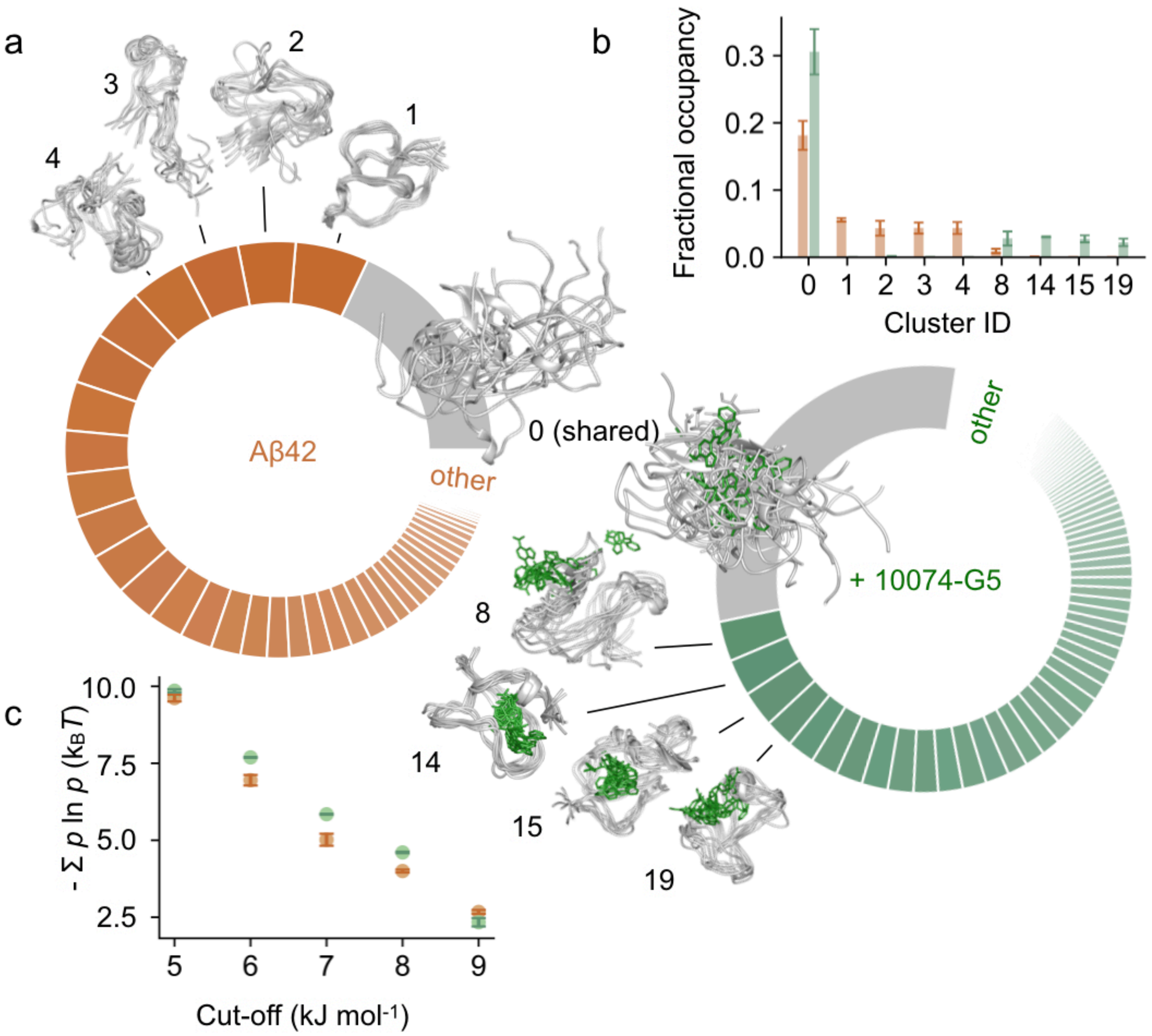


**Figure S7. Clustering of the conformational states in the unbound and bound structural ensembles of Aβ42. (a)** Donut plots quantifying the conformational states of Aβ42 in the unbound (left, orange) and bound (right, green) simulations. Clustering was performed on concatenated trajectories by considering only Aβ42. Inter-residue contact maps based on the Lennard-Jones potential were used as input for the GROMOS clustering algorithm (see **SI**). The data shown is for the cut-off value of 8.5 kJ mol$^{-1}$. Each slice represents a distinct state. The simulations share one major state (Cluster 0, grey), which comprises 18 and 31% of the unbound and bound ensembles, respectively. **(b)** Convergence of the 5 most populated clusters for the Aβ42 ensembles in the absence and presence of 10074-G5. Bar plot shows fractional cluster occupancies for the five most populated clusters for the unbound (orange) and bound (green, right) simulations. Frames from approximately the first 14 μs of equilibration were discarded. Fractional cluster occupancies were calculated on 35,000 frames for each concatenated trajectory sampled based on metadynamics weights. Error bars represent standard deviations between fractional occupancies calculated on the first and second halves of the analysed trajectories. **(c)** The conformational entropy of Aβ42, estimated via Gibbs entropy, is consistently higher in the 10074-G5-bound form of the peptide for several clustering cut-off values. The conformational entropy was calculated such that the weights, $p$, of each state correspond to the fractional occupancy as determined by the GROMOS clustering algorithm (see **SI**). Error bars represent $\pm$ standard deviations of the conformational entropy calculated from the first and second halves of the simulations.

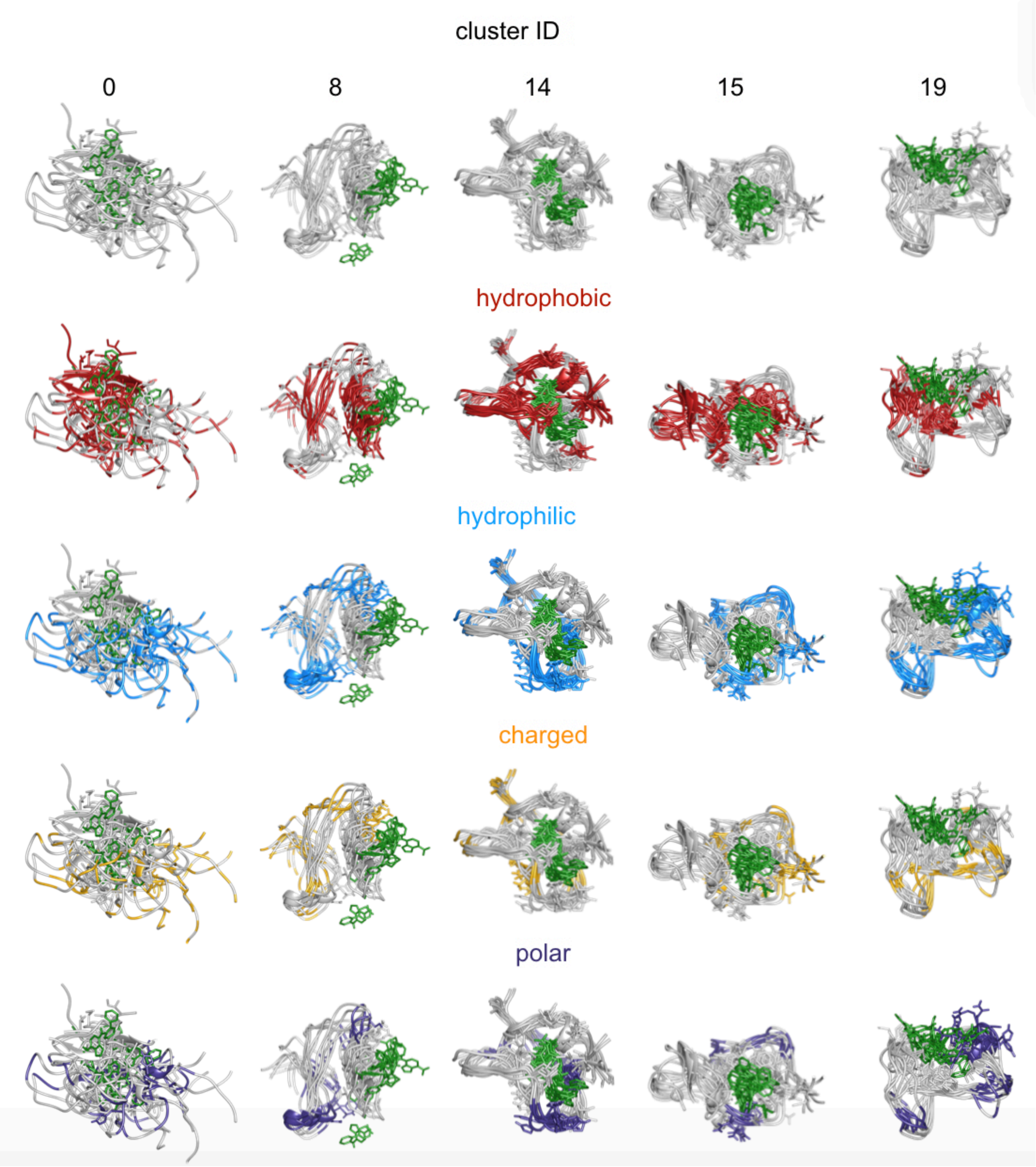


**Figure S8. Illustration depicting key groups of residues for heterogeneous bound clusters of Aβ42.** Depiction of several conformations of Aβ42 (grey) with 10074-G5 (green) showing that many types of residues are involved in the binding interaction. Hydrophobic residues (red) include Ala, Val, Ile, Leu, Met, Phe, and Tyr. Hydrophilic residues (blue) His, Ser, Asn, Gln, Tyr, Arg, Asp, Lys, and Glu. Charged residues (yellow) include Arg, Asp, Lys, and Glu. Polar residues (indigo) include His, Ser, Asn, Gln, and Tyr.

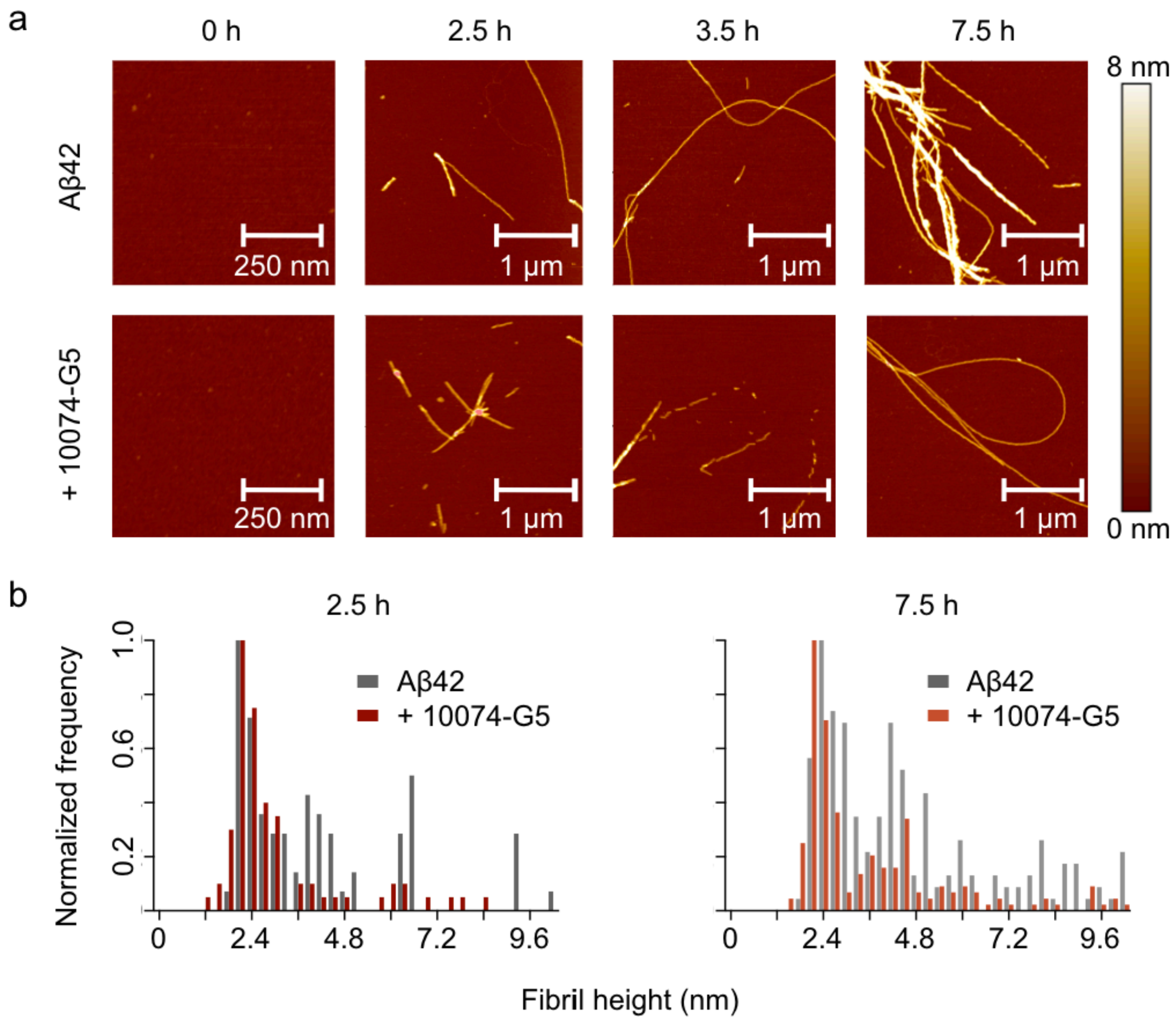


**Figure S9. AFM measurements show that 10074-G5 slows the formation of** Aβ42 **aggregation**. **(a)** Representative 3-D morphology maps from AFM time course experiments showing fibrils formed from 1 μM Aβ42 in the presence and absence of 6 μM 10074-G5. **(b)** Single-molecule statistical analyses of maps such as those shown in (a). Bar plots show distributions of the cross-sectional heights of the Aβ42 fibrils at 2.5 (left, N=75 per condition) and 7.5 h (right, N=200 per condition) in the presence and absence of 10074-G5. These data demonstrate that in the absence and presence of 10074-G5, Aβ42 forms protofibrillar aggregates with average cross-sectional diameter of approximately 2-3 nm and mature fibrillar aggregates with average diameter of approximately 5-6 nm for fibrils formed collected in the presence and absence of 10074-G5. Notably, however, at the same timepoint, fibrillar aggregates formed in the presence of 10074-G5 have smaller cross-sectional diameters than those formed in its absence, with a significantly higher proportion of protofibrillar species as compared to mature fibrillar species. These data suggest that fibril formation and maturation of cross-β sheet structure is retarded by 10074-G5.

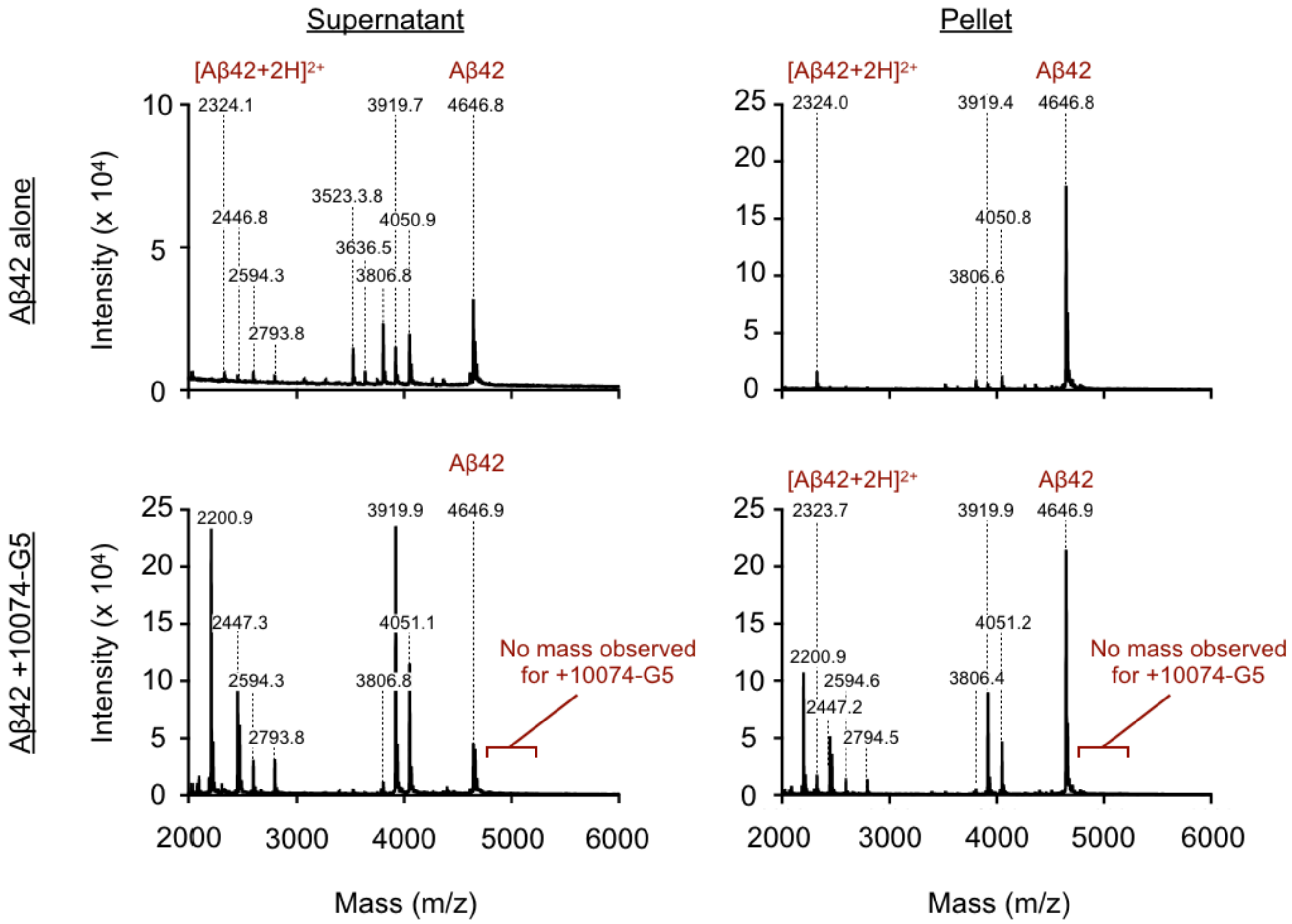


**Figure S10. Matrix assisted laser desorption/ionization (MALDI) mass spectrometry of Aβ42 in the presence and absence of 10074-G5.** 15 µM of monomeric Aβ42 was aggregated overnight at 37 °C in the absence (top) and presence (bottom) of 10074-G5. Samples were spun down to separate the supernatant (left) and pellet (right) (see **Materials and Methods**). MALDI mass spectrometry shows no chemical modification of Aβ42 by the presence of 10074-G5.

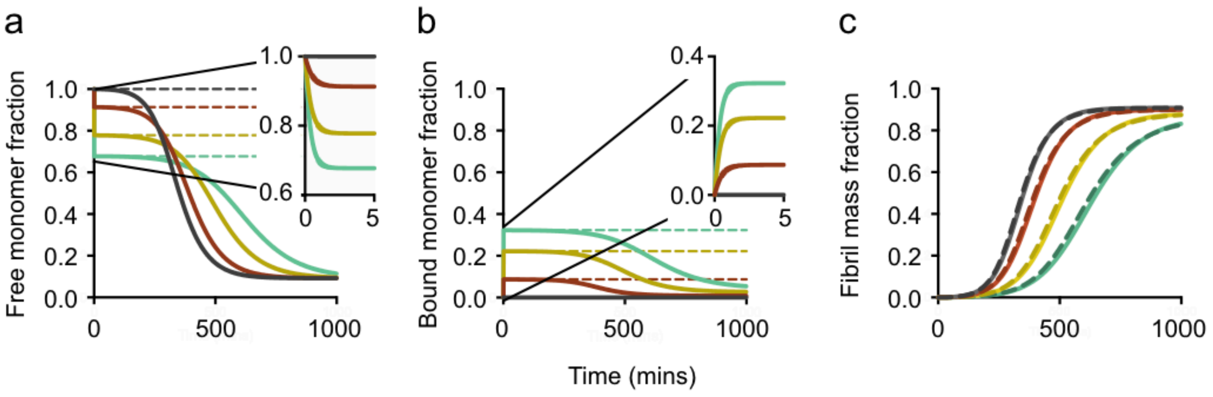


**Figure S11. Behaviour of solution to monomer sequestration model (Eq S11). (a)** The time evolution of the free monomer concentration shows an initial rapid phase of pre-equilibration to Eq. S12 (dashed lines), followed by slower aggregation ending at the critical concentration. **(b)** The time evolution of the concentration of bound monomers also displays two phases: a rapid pre-equilibration to Eq. S12 and slow aggregation ending at Eq. S13. **(c)** Time evolution of aggregate mass concentration is described by Eq. S10 with effective parameters from Eq. S14. A comparison between analytical (dashed lines) and numerical (solid lines) solution to the monomer sequestration model, Eq. S11, shows excellent agreement.

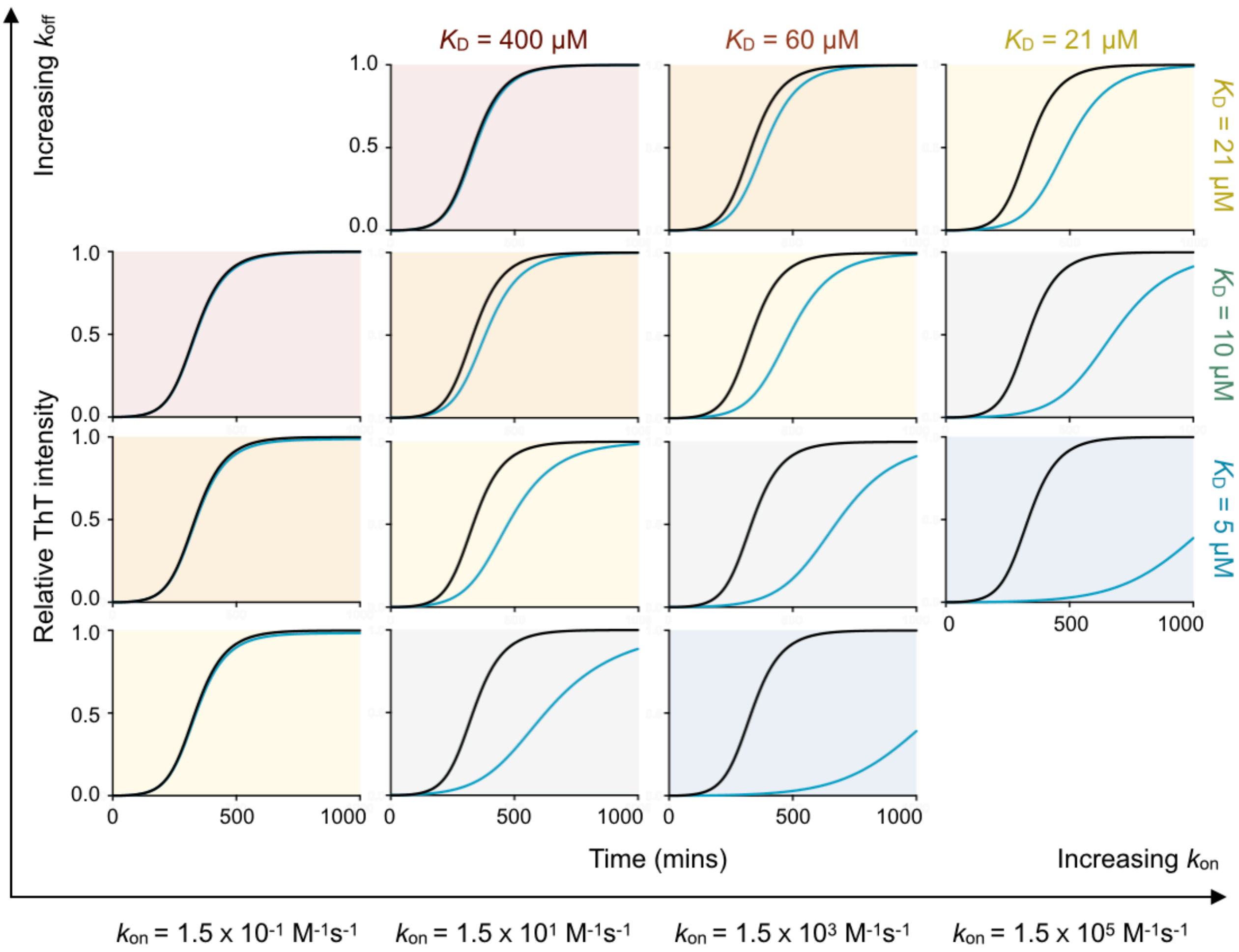


**Figure S12. Phase diagram illustrating numerical solutions to the kinetic equations for different $k_{on}$ and $k_{off}$ rates.** Curves represent kinetic aggregation traces of 1 µM Aβ42 in the absence (black) and presence of 6 µM of compound (blue). Diagonals correspond to constant values of $K_D = k_{off}/k_{on}$ . The values of $k_+k_n$ and $k_2k_+$ are the same as those shown in **Figure 4a** of the main text.

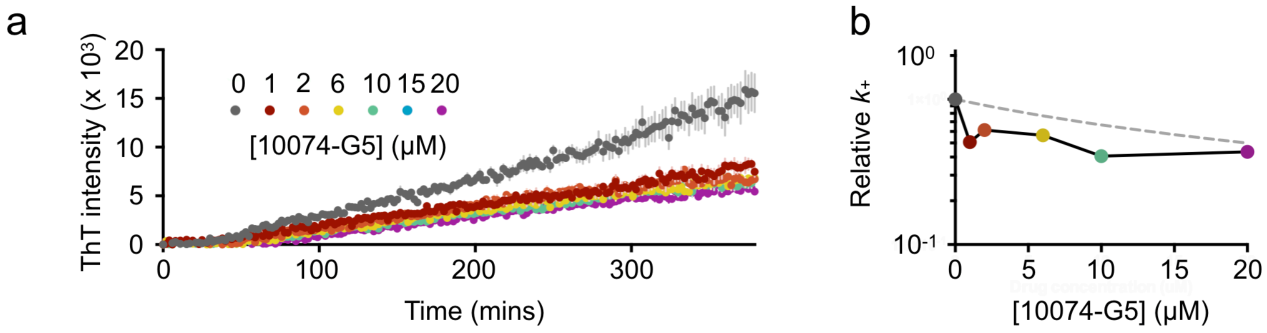


**Figure S13. Measurement of the effects of 10074-G5 on the fibril elongation rate. (a)** ThT aggregation experiments performed in the presence of 15% preformed fibrils in the presence and absence of varying concentrations of 10074-G5. Error bars represent ± one standard deviation. Measurements were taken in triplicate. **(b)** Change of the elongation constant, $k_+$, as a function of the concentration of 10074-G5, determined from the data shown in panel (a). Grey dashed line represents the theoretical prediction of effective elongation rates predicted from Eq. S14c.

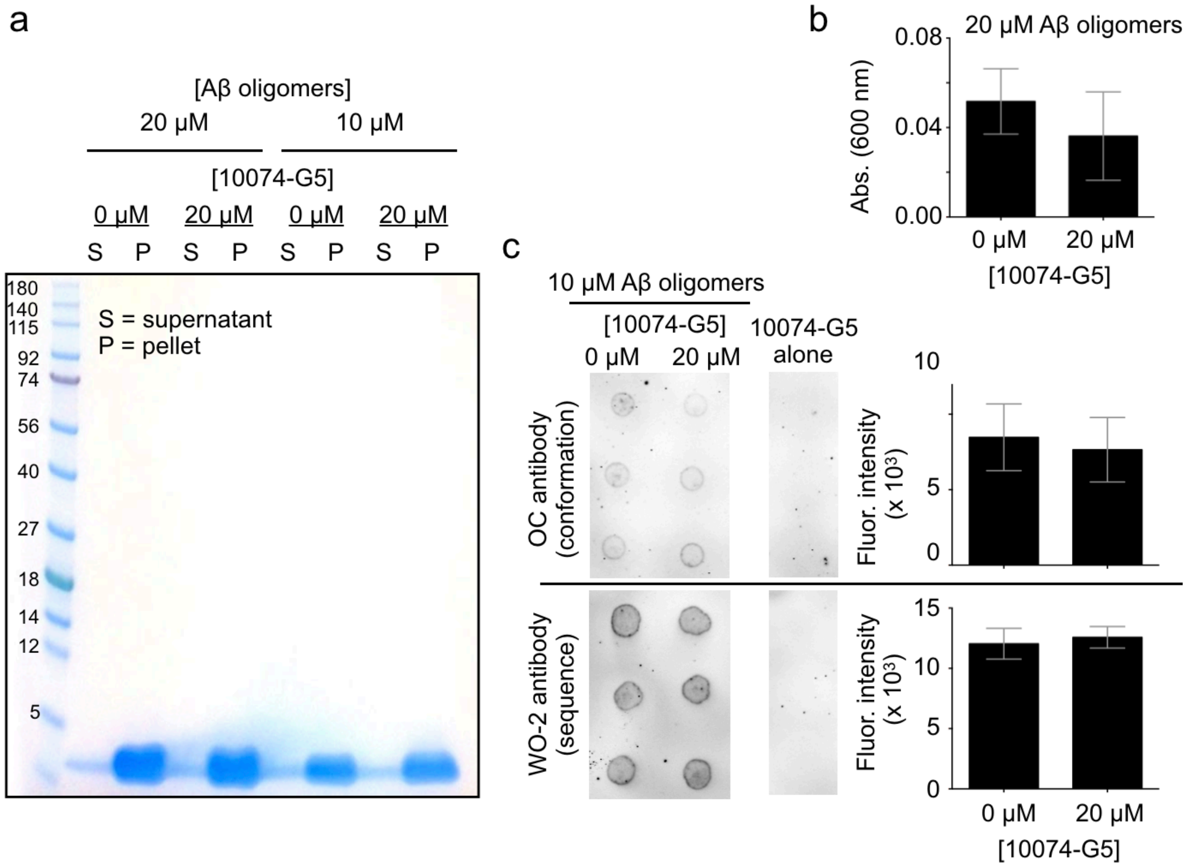


**Figure S14. 10074-G5 does not alter pre-formed Aβ40 oligomers. (a)** SDS gel of samples containing pre-formed $Zn^{2+}$-stabilised Aβ40 oligomers (20 μM and 10 μM) after incubation in the presence and absence of 20 μM 10074-G5. Samples were centrifuged to identify whether oligomers were dissociated into monomer upon addition of the compound. These data demonstrate that the compound does not dissociate pre-formed oligomers. **(b)** Turbidimetry measurements of 20 μM pre-formed Aβ40 oligomers show no significant difference in the presence of the compound, suggesting that 10074-G5 causes neither dissociation nor clustering of these oligomers. Measurements were background subtracted from buffers in the presence and absence of the compound, respectively. Error bars represent ± one standard deviation. Measurements were taken in triplicate. **(c)** Dot blot using the OC-antibody shows that pre-formed oligomers maintain their cross β-sheet content in the presence of the 10074-G5. Error bars represent ± one standard deviation. Measurements were taken in triplicate, as shown.

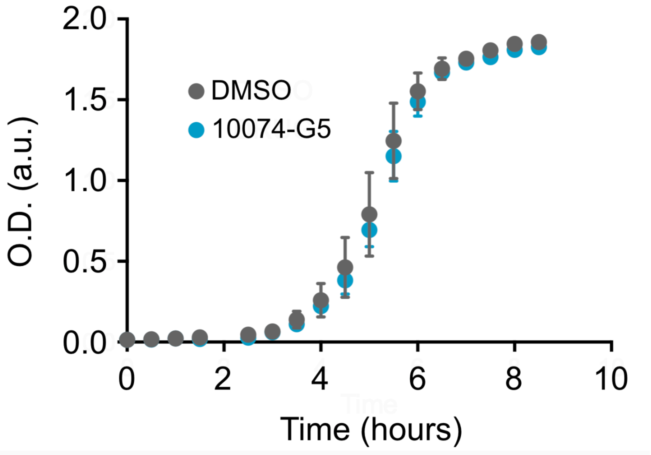


**Figure S15. 10074-G5 does not alter the growth of *E. coli* strain OP50.** Optical density (O.D.) growth curves of *E. coli* strain OP50 in the presence (blue) and absence of (grey) of 5 µM 10074-G5, showing that 10074-G5 does not alter the *E. coli* consumed by the *C. elegans*.

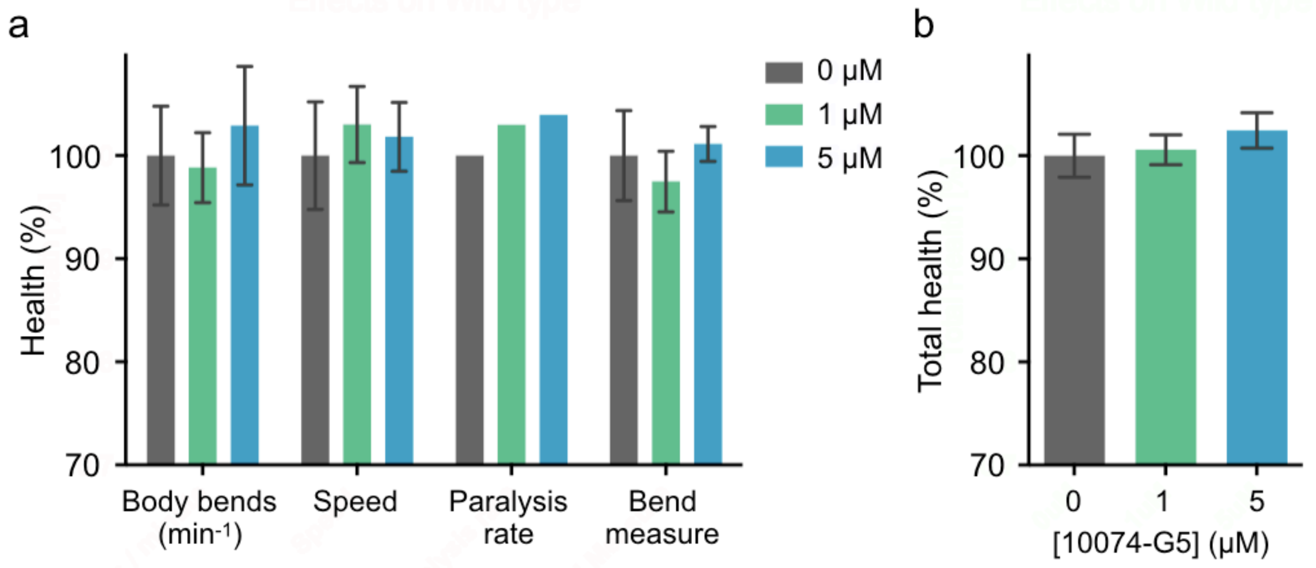


**Figure S16. Effects of 10074-G5 on wild-type *C. elegans*. (a)** Health scores (%) for the rate of body bends, the speed of movement, and the paralysis rate, and the magnitude of body bends at day 6 of adulthood. Error bars represent ± SEM, N=150. **(b)** Combined total health scores from panel (a). Error bars represent ± SEM.
